# Unveiling protist composition and diversity patterns with eDNA metabarcoding: comparing short- and long-read approaches

**DOI:** 10.64898/2026.02.07.704525

**Authors:** Skouroliakou Dimitra-Ioli, Dupont Deborah Whitney Euselia, Vandenboer Yelle, D’hont Sofie, Sabbe Koen, Schön Isa

## Abstract

Environmental DNA (eDNA) metabarcoding is a key tool in biodiversity monitoring due to its high-throughput, non-destructive nature. While short-read (SR) sequencing platforms such as Illumina Miseq have been routinely used in environmental monitoring, their limited read lengths (less than 600 bp) constrain the depth of taxonomic assignment, particularly for complex microbial eukaryotes like protists. Conversely, long-read (LR) sequencing technologies like Oxford Nanopore Technologies (ONT) offer promising alternatives but remain underutilized for studying protist communities. We conducted a comparative study of SR versus LR metabarcoding of protist communities along a coastal-offshore gradient in the Belgian part of the North Sea. Using amplicons targeting the V4 region (SR; 577 bp) and the V4–V5 region (LR; 745 bp) of the 18S rRNA gene, we compared diversity patterns, taxonomic assignment, and community composition between approaches. We observed general congruence in community composition at higher taxonomic levels, but under the applied workflows, LR metabarcoding yielded a greater depth of taxonomic annotation at lower taxonomic ranks. Notably, dinoflagellates were less overrepresented in LR data, and a unique detection of potential nuisance taxa (e.g., *Bellerochea*), and ecologically important genera such as haptophytes (e.g., *Gephyrocapsa*) was achieved. These results highlight the potential of LR metabarcoding to complement SR approaches by providing increased taxonomic annotation depth and ecological insights. Although both methods targeted only partial regions of the 18S rRNA gene, LR metabarcoding yielded a greater depth of taxonomic assignment under the applied workflows. As next-generation sequencing technologies continue to evolve, our research provides valuable insights for selecting optimal strategies in routine plankton monitoring and biodiversity assessment programs.

## Introduction

Over the past decade, eDNA metabarcoding has started to transform monitoring as a relatively fast and non-destructive method to characterize biodiversity and its changes worldwide (Ruppert *et al*., 2019). Environmental DNA (eDNA) refers to the genetic material that can be extracted from environmental samples of different matrices. It is acknowledged that eDNA contains a complex mixture of intracellular DNA, originating from living cells or potentially whole organisms, and extracellular DNA resulting from natural cell death and/or destruction of cell structure (Taberlet *et al*., 2012; Pawlowski *et al*., 2020; Rodrigez-Espeleta *et al*., 2021). Amplicon sequencing of eDNA facilitates identification of the composition of natural assemblages without prior knowledge and also detects small-size organisms like pico- and nanoplankton, cryptic species with undistinguishable morphology, and rare species (López-García *et al*., 2001; Sunagawa *et al*., 2015; Burki *et al*., 2021; Duarte *et al*., 2023). It has been routinely used to study protist communities in various coastal systems (e.g., Lambert *et al*., 2019; Caracciolo *et al*., 2022; Longobardi *et al*., 2022; Skouroliakou *et al*., 2022). As its use becomes more widespread, eDNA is increasingly integrated into international monitoring initiatives aimed at evaluating the environmental status of marine ecosystems and predicting future changes, such as the European Marine Omics Biodiversity Observation Network (EMO BON) (Cardinale *et al*., 2012; Santi *et al*., 2023).

Protists form a significant part (over 60%) of marine biomass (Bar-On and Milo, 2019) playing a key role in biogeochemical cycles and carbon sequestration and as primary producers in marine food webs (Falkowski *et al*., 1998; Field *et al*., 1998). Investigations of protists with eDNA have so far been based on short-read sequencing methods provided by Illumina technology (named short-read (SR) metabarcoding hereafter), targeting hypervariable parts of the 18S ribosomal region such as V4 (Pernice *et al*., 2013), often in conjunction with V5 or V9 (Amaral-Zettler *et al*., 2009). This approach has been able to identify diversity patterns of protist communities and their major drivers. For example, SR metabarcoding data revealed correlations of seasonal and/or spatial variation in protist diversity with environmental fluctuations (Caracciolo *et al*., 2022), interspecific interactions (Genitsaris *et al*., 2015), and water depth (e.g. Meziti *et al*., (2023). Protist species causing harmful algal blooms (HABs) have also been successfully detected with SR metabarcoding at low cell concentrations; an example is the HAB of *Lepidodinium chloroforum* that was identified outside of algae blooming periods at the coast of south Brittany (Roux *et al*., 2023)

Despite its widespread application, short-read (SR) metabarcoding provides limited molecular information because commonly sequenced amplicons are typically short (approximately 300 to <600 base pairs). This restricted sequence length can lead to biased estimates of taxonomic composition, resulting in the over- or underrepresentation of certain taxa. For example, dinoflagellates, which possess numerous ribosomal gene copies per cell, are frequently overrepresented due to PCR and sequencing biases (Prokopowich et al., 2003; Wisecaver and Hackett, 2011). In addition, SR metabarcoding often limits taxonomic identification to higher taxonomic ranks (Szoboszlay et al., 2023).

These limitations are particularly pronounced for protists, which comprise deeply divergent evolutionary lineages and exhibit extensive cryptic diversity as well as highly variable rates of rRNA gene evolution across taxa (Burki et al., 2021; Jamy et al., 2022). Accurate characterization of protist communities therefore requires sufficient phylogenetic signal to discriminate among closely related taxa and evolutionary lineages. Although SR metabarcoding has proven effective for detecting broad diversity patterns and ecological gradients in protist communities (e.g., Genitsaris et al., 2015; Caracciolo et al., 2022; Meziti et al., 2024), the limited length of commonly used markers constrains the depth of taxonomic assignment achievable for many groups—particularly those that are underrepresented or unevenly annotated in reference databases (Szoboszlay et al., 2023). In contrast, longer amplicons provide increased sequence context and, in principle, greater potential for improved taxonomic resolution and discrimination among closely related protist taxa (Jamy et al., 2022; Gaonkar et al., 2024; Chwalińska et al., 2025).

Long-read sequencing technologies such as Oxford Nanopore Technologies (ONT) have been developed in recent years. ONT long-read metabarcoding has successfully characterized bacterial (Stoeck *et al*., 2024; Van der Loos *et al*., 2021), and zooplankton communities (Semmouri *et al*., 2021) with eDNA. However, to our knowledge, only few studies have compared protist diversity and composition between SR and LR metabarcoding approaches based on 18S rDNA (Jamy et al., 2022; Gaonkar et al., 2024; Chwalińska et al., 2025). Few other studies focused on specific protist groups such as Radiolaria (Sandin *et al*., 2022).

Long Read metabarcoding is advantageous in providing more DNA sequencing data from longer amplicons of the target DNA regions (more than 600 bp) and thus could potentially enable deeper taxonomic annotation and discrimination among closely related taxa. This could be particularly beneficial for taxonomically diverse groups like protists, where fine distinctions between closely related taxa are often required for accurate ecological assessments (Gaonkar and Campbell, 2024). The most significant limitation of ONT, however, is its greater error rate as compared to Illumina sequencing based on base calling biases (Wang *et al*., 2021). Given the recent improvements in the ONT library kits and flow cell chemistry both reducing sequencing errors (Wang *et al*., 2021), it has now become possible to test whether LR metabarcoding using ONT can overcome the limitations of SR metabarcoding based on Illumina sequencing. As a result, differences between SR and LR outputs may reflect both biological signal and methodological artefacts. Evaluating how these approaches complement each other, and to what extent they can be integrated in comparative ecological analyses, remains an open question for protist metabarcoding.

The present study focuses on protist communities, comparing their taxonomic assignment, composition, and diversity across a coastal-offshore gradient between metabarcoding data generated with (SR) and (LR) sequencing. We expect congruence in diversity patterns based on higher taxonomic levels between both methods similarly to Gaonkar and Campbell (2024). Because LR metabarcoding generates longer sequence reads with increased phylogenetic information content, we hypothesize that LR metabarcoding will provide greater depth of taxonomic assignment for protist communities under the applied workflows, potentially revealing a higher number of taxa at lower taxonomic ranks compared to SR approaches (Jamy et al., 2022; Gaonkar and Campbell, 2024; Chwalińska et al., 2025). In the context of the increasing integration of eDNA metabarcoding into studies of biodiversity, ecology and also monitoring, such comparative studies are urgently required to inform about methodological choices. Our work contributes practical insights into the strengths and limitations of commonly used sequencing strategies for the characterization of marine protist communities.

## Material and Methods

### Study area

The Belgian Part of the North Sea (BPNS; 3,454 km^2^) is an epicontinental shallow area (maximum 35 m deep) with a coastal length at ca. 66 km (Mortelmans *et al*., 2019). The BPNS is affected by Atlantic waters through the English Channel and by freshwater inputs from the Scheldt and Meuse rivers (e.g., Aubert *et al*., 2022 and references therein). It is macrotidal and well-mixed without seasonal stratification (e.g., Blauw *et al*., 2012). The BPNS experienced eutrophication (1950s–1980s), de-eutrophication (1980s–2000s), and an increase in water temperature since the 1970s (Beaugrand, 2004). This area is furthermore heavily impacted by the introduction of non-indigenous species, industrial and agricultural pollution, overfishing and trawling, offshore wind farming and heavy shipping traffic (Emeis *et al*., 2015). The protist communities of the BPNS have been extensively studied in the past with a wide range of molecular and classic microscopic and imaging techniques (e.g., Breton *et al*., 2006; Nohe *et al*., 2020; Aubert *et al*., 2022; Perneel *et al*., 2024), making it an ideal area for the current study.

### Sample collection

A total of 79 samples were collected aboard the *R*.*V. Belgica* during 15 campaigns at a monthly basis during the regular monitoring program organised by RBINS (Table S1). Three stations were sampled from February 2022 to May 2023; the coastal station MOW1 (51° 21.50’ N, 3° 07.50’ E), the transitional station WO5 (51° 25.00’ N, 2° 48.50’ E), and the offshore station WO8 (51° 27.61’ N, 2° 20.91’ E) situated 87 km west of the coastal station (Figure S1). No samples were collected in May, November 2022, and January 2023 due to rough weather conditions and/or the boat’s unavailability. At each station, sea water was collected with Niskin bottles at 1 m depth (subsurface) and 1 m above the seabed (epibenthic). Seawater ranging in volume from 0.1 to 1 L was filtered on board with a low-pressure vacuum pump through 0.45□μm membrane filters (47□mm, Merck-Millipore), depending on the quantity of suspended matter in the water (i.e., until clogging of the filter occurred). All filters were stored at −80 °C onboard the vessel, after snap-freezing in liquid nitrogen, until DNA extractions took place in the dedicated eDNA laboratory at the RBINS. Amplification and sequencing procedures used short-read (MiSeq, Illumina) and long-read (GridIon, Oxford Nanopore Technologies) sequencing approaches (Figure S2). Here, 66 subsurface and 13 epibenthic water samples were analysed together with five field and three laboratory negative controls to assess potential contamination.

### DNA extraction

DNA was extracted from half of the filters following an adapted protocol of the DNeasy PowerLyzer Microbial kit (Qiagen, Germany). Modifications were made to maximise the yield, such as increasing the rotation speed of the PowerLyzer (30 Hz for 3 minutes), reducing the volume of eluents to 35 µl and repeating elutions three times.

### Primer selection and in silico tests

One of the most widely used eukaryote-specific primer pairs in SR metabarcoding studies are the forward primer TAReuk454FWD1 (CCAGCASCYGCGGTAATTCC) and the reverse primer TAReukREV3 (ACTTTCGTTCTTGATYRA) (Stoeck *et al*., 2010), which together amplify 577bp of the V4 region of the 18S rRNA gene. We used the same primers for LR metabarcoding as Semmouri *et al*. (2021), who successfully characterized marine zooplankton communities of the North Sea in high resolution. The 18S primers F-566 (CAGCAGCCGCGGTAATTCC) and R-1200 (CCCGTGTTGAGTCAAATTAAGC) (Hadziavdic *et al*., 2014) amplify the V4 and V5 regions of the 18S rRNA gene with a total length of 745 bp (Figure S3). To assess how well these primers would amplify protists from a reference database, *in silico* tests were performed first using the PR2 Primer Database v.2 (Vaulot *et al*., 2022), which is curated to the species level for protists. Primers were matched with all protist sequences available in the database, allowing to identify mismatches and evaluate amplification efficiency. The SR primer pair TAReuk454FWD1 **–** TAReukREV3 (Stoeck *et al*., 2010) detected in silico 83.2% of all eukaryotic barcodes in the PR2 database; and the LR primer pair F-566 **–** R-1200 (Hadziavdic *et al*., 2014) 90.45% (Figure S4 and S5).

### Short-read amplification and library preparation

Forward and reverse primers with overhang and adapter sequences of 38 bp were used, which bind to Illumina indexes and sequencing adapters according to the standardised Illumina protocol. Samples were amplified in 25μl reactions using the Kapa HotStart ReadyMix DNA polymerase (Roche Sequencing Store) with the following PCR settings: 95°C for 3 min, 30-35 cycles (depending on the amount of DNA) including 95°C for 30 sec, 58-52°C for 30 sec, 72°C for 1 min, and a final elongation step of 5 min at 72°C. All PCRs were conducted in replicates. Successful amplification was confirmed on 1% agarose gels. PCR products were purified with the AMPure XP bead-based reagent purification kit (Beckman Coulter Life Sciences). After the first amplification with overhangs had been performed, dual indexes barcodes (Nextera XT index kit v2 Set A, B, C) were ligated to the adapter sequences allowing multiplexing of PCR products from different samples. For the barcoding PCRs, the KAPA Hotstart ReadyMix DNA polymerase was also used in final volumes of 50µl. PCR settings were: 95°C for 3 min, 8 cycles at 95°C for 30 sec, 55°C for 30 sec, 72°C for 30 sec, and a final elongation step of 5 min at 72°C. PCR products from each sample were pooled and purified using the AMPure XP bead-based reagent purification kit. DNA concentrations after barcoding and purification were measured with a Qubit dsDNA BR assay (ThermoFisher) and adjusted to 45 nM.

### Short-read sequencing and bioinformatic analysis

Pooled and purified amplicons were sequenced in paired-end mode on an Illumina MiSeq 2□×□300 platform (Genewiz, Germany GmbH, Leipzig). Quality filtering of reads, identification of amplicon sequencing variants (ASVs), and taxonomic affiliations were conducted with the R-package *DADA2* v.1.26.0 (Callahan *et al*., 2016). Due to the poor quality of reverse sequences (Figure S6A), only the forward reads were retained for subsequent bioinformatic analysis (Figure S6B). A total of approximately 17,000,000 high quality forward reads were obtained, while only 8,000 reads were recovered when using paired end reads (forward and reverse) (Figure S7). The final dataset with forward reads only included 87 samples (79 field samples, five field controls, and three lab controls). Forward reads were trimmed at position 280, primers were removed (*TrimLeft*) and reads with ambiguous nucleotides or with a maximum number of expected errors (*maxEE*) exceeding 2 were filtered out using the function *filterAndTrim()*. Chimeric sequences were identified and removed using DADA2’s consensus-based chimera removal approach (*BimeraDenovo, pooled method*). To assign taxonomy to ASVs, the default RDP naive Bayesian classifier method was used with PR2 v5.0.0. *(assignTaxonomy)*, including a minimum bootstrap confidence threshold (minBoot = 50). Under this setting, taxonomic assignments were truncated when bootstrap support fell below the threshold. At the end of the analyses, a total of 12,627,934 eukaryotic reads and 17,543 ASVs were obtained.

### Long-read amplification and library preparation

Samples were amplified with the LR primer pair F-566 **–** R-1200 in 25μl reaction volumes using the KAPA HotStart ReadyMix DNA polymerase (Roche Sequencing) and the following PCR protocol: 94°C for 50 sec, 25–30 cycles including 94°C for 50 sec, 63°C for 50 sec, 72°C for 1 min, and a final elongation step of 10 min at 72°C. Successful amplification was confirmed by 1% agarose gel electrophoresis with subsequent staining with Midori Green. Amplicons from single PCR reactions were purified with AMPure XP (Beckman Coulter Life Sciences) before LR sequencing at OHMX.bio (Ghent, Belgium). The libraries were prepared with the Native Barcoding kit 96 V14 (SQK-NBD114.96, ONT) and the manufacturer’s protocol. Uniquely barcoded DNA amplicons were pooled, purified (0.4X AMPureXP beads), adapter-ligated, and repurified. Libraries were quantified (Qubit dsDNA HS assay, ThermoFisher) before adding Sequencing Buffer (SB) and Loading Beads for sequencing.

### Long-read sequencing and bioinformatic analysis

Sequencing was performed on a GridION R10.4 flow cell, preliminarily primed with Flow Cell Flush (FCF), Bovine Serum Albumin (BSA), and Flow Cell Tether (FCT) as priming mix. Twenty femtomoles of pooled libraries were loaded three times through the SpotON sample port and sequenced for 72 hours with MinKNOW high-accuracy base calling; altogether, 22.5 million raw long-reads were generated. The quality of the raw sequencing reads was controlled with PycoQC v.2.5.2, in which sequences were basecalled, demultiplexed with Guppy v6.4.8 (Nanopore) and low-quality LR data were removed with the >PHRED9 parameter. Then, the quality of demultiplexed FASTQ files was visualized with *NanoPlot* v1.46.2 (De Coster *et al*., 2023). Based on the primer pair, the expected long-read amplicon size was approximately 750 bp. Long-reads were filtered using a length range of 600–1000 bp to retain target amplicons, while allowing for expected variation in amplicon length. The resulting read-length distribution corresponded to the expected amplicon size, with a median read length of 762 bp (Figure S9).Therefore, long-reads ranging between 600 and 1000 bp were kept with the *chopper* tool v0.12.05 and with read quality scores exceeding the median score value of the entire LR dataset (i.e., q=14; Figure S8) (De Coster *et al*., 2023). Profiling was done with EMU v3.5.0 (Expectation-Maximization algorithm) with the *keep-counts* command, and the PR2 v5.0.0_emu reference database. Filtered long reads were aligned to the PR2 reference sequences using *minimap2* v2.24, as implemented within the EMU pipeline. This pipeline is designed to estimate relative abundances at the species-level in two steps: first, it aligns reads to a reference database, and second, it applies an expectation-maximization-based error correction. Specifically, it assigns relative abundances by computing the likelihood that each read comes from each species using probabilistic alignment and iterating to refine initial estimates. This method improves the accuracy of community profiles at the genus and species levels, particularly with error-prone reads (Curry *et al*., 2022). Our LR approach generated a total of 13,898,821 long reads and identified 1,752 species.

### Taxonomic filtering in protist metabarcoding datasets

Samples from the negative controls were excluded from the downstream analysis of both SR and LR metabarcoding datasets. For the SR metabarcoding dataset, all amplicon sequence variants (ASVs) were pooled to the species level using the *tax_glom* function *in phyloseq*; (McMurdie and Holmes, 2013) to facilitate comparisons with the LR dataset. Given that the focus of this study was on protists, the following taxonomic groups were retained: Amoebozoa, Archaeplastida, CRUMs, Excavata, Haptista, TSAR, and Cryptista kingdoms. Within the Obazoa kingdom, the Breviatea division and the Ancyromonadida, Nibbleridia, and Apusomonada supergroups were included, as well as the genus *Tunicaraptor*. In the Opisthokonta domain, the Choanoflagellata division was considered (Figure S10). Sequences associated to the domains Bacteria, Fungi, Metazoans, the Ichthyosporea, the Streptophyta supergroup, as well as sequences corresponding to chloroplasts, mitochondria and any unclassified taxa at the domain level were excluded to ensure accurate analyses of protist data. Additionally, unassigned taxa at the species level were also removed. In total, 7,438,382 reads were identified as protists and classified as 1,039 species based on the SR metabarcoding dataset, while 7,623,617 reads were identified as protists in 1,180 species from the LR metabarcoding dataset. The relative abundance of protist taxa was calculated for each dataset including the taxonomic groups described above.

All bioinformatic analyses were conducted on a local server to increase computational efficiency.

### Statistical analysis

Visualization of data and statistical analyses were performed in R version 4.1.0. (R Core Team, 2021) using the *phyloseq* (McMurdie and Holmes, 2013), the *ggplot2*, and the *vegan* packages (Wickham, 2016). As eDNA metabarcoding data are compositional and influenced by sequencing depth, read counts were normalised across samples prior to alpha diversity analysis. For this purpose, samples within each dataset were rarefied to the lowest sequencing depth observed (10,131 reads for both datasets). Rarefaction is commonly applied in metabarcoding studies to reduce biases associated with uneven sequencing effort (Schloss et al., 2024), and unequal sampling effort (e.g., Bruce et al., 2021; Ramond et al., 2021). To avoid any further bias by unequal sample size, Random Under-Sampling (RUS) was applied using the base R sample function (Dittman et al., 2014). After rarefaction and resampling, both SR and LR datasets contained 76 samples. Alpha diversity indexes (richness, Shannon, Gini-Simpson (1-D), and Chao1) were calculated to describe and compare diversity of protist communities. The Kruskal-Wallis test was used to evaluate if protist alpha diversity significantly differed among stations, and the Wilcoxon test was applied to test for differences in alpha diversity between metabarcoding methods. To further describe protist patterns, three thresholds for rare taxa were defined for each dataset (SR and LR datasets, respectively): *i*) species were considered as abundant if they were present in more than 0.1% of the average relative read abundance, *ii*) rare species were present in less than 0.001% of the average relative read abundance, and *iii*) intermediate rare species contributed to 0.001 and 0.1% of the average read relative abundance (Logares *et al*., 2014). NMDS ordination plots with a centred log-ratio (CLR) transformation were constructed to evaluate whether protist communities were structured according to the coastal-offshore gradient; NMDS plots used Bray-Curtis similarities based on a non-transformed number of rarefied reads for each dataset. Then, PERMANOVA analyses were used to investigate what proportion of the variance in community composition was explained by the location (Anderson, 2017).

## Results

### Numbers of sequencing read and taxonomic assignment of SR and LR metabarcoding

Potential contamination was assessed first from negative field and lab controls. Protist sequences detected in the field samples were genuine and not the result of contamination, given that field samples clearly contained much higher frequencies of protist taxa (98% for SR and 85% for LR metabarcoding) than the negative controls (Figure S11).

A total of 17 million eukaryotic reads were generated using SR Illumina sequencing, whereas LR Nanopore sequencing produced 22 million reads. After removing primers and filtering, approximately 14 million reads were retained for the LR, and 13 million for the SR dataset for the 79 field samples. Both sequencing platforms detected a similar number of reads being assigned to protists. SR produced a total of 7,438,382 reads identifying 1,039 species, and the LR metabarcoding dataset resulted in 7,623,617 protist reads, with 1,180 identified species. In terms of relative abundance of total reads (%) for each dataset, protists contributed more than 60% to read relative abundance in both datasets, with SR exhibiting a 10% higher relative abundance of reads as compared to LR (Figure S12). The taxonomic assignment of protists varied greatly between the two sequencing methods. The LR metabarcoding dataset contained only 1.4% of reads being unassigned to protists at the species level and 0% at the genus level. In contrast, the SR metabarcoding dataset had a ten times higher proportion of unassigned reads, with 12% being unassigned at the genus level and 15.2% at the species level (Figure S13).

### Protist composition across taxonomic levels

With both metabarcoding techniques, TSAR (Telonemia, Stramenopiles, Alveolata, and Rhizaria) were the most prevalent kingdoms, being identified as 87.4% of relative read abundances in the LR and 81.7% in the SR dataset (Figure 1A), respectively. The second most dominant kingdom, Haptista, contributed 4.2% of relative read abundances in the LR and 7.2% in the SR dataset. At the supergroup level, Stramenopiles and Alveolata were the most dominant groups. Stramenopiles dominated the LR dataset with 43.5%, as compared to 33.2% in the SR dataset (Figure 1B). In contrast, Alveolata was with 38.4% more abundant in the SR dataset as compared to 35.9% in the LR dataset. This pattern was similar at lower taxonomic levels (Figure 1C–H). Specifically, Gyrista was the most abundant division in LR (40.2%), while in SR, Gyrista (29.5%) and Dinoflagellata (28.5%) were nearly equally represented (Figure 1C). At the class level (Figure 1D), Mediophyceae prevailed in the LR (24.5%) and Dinophyceae in the SR sequencing reads (19.2%). Spirotricheae were present in both datasets in comparable read relative abundances (LR: 9.5%, SR: 8.4%) while Syndiniales, Prymnesiophyceae, and Filosa-Thecofilosea were slightly more abundant with SR. The order Thalassiosirales dominated in LR (12 %) and Gymnodiniales in SR (10.4%) datasets, respectively (Figure 1E). Similarly, at the family level, Thalassiosiraceae (7.8%) and Gymnodiniaceae (8.6%) prevailed in the LR and SR results, respectively (Figure 1F). The genus *Thalassiosira* was more prevalent in the LR data set (7.5%), while *Gyrodinium* was more abundant in the SR dataset (Figure 2G). It is worth noting that the SR dataset did not detect the same dominant diatom genera as LR: *Chaetoceros* (5.6%) was most frequent with SR, while *Thalassiosira* (4.1%) was dominating with LR. The species *Thalassiosira curviseriata* was dominant with LR (4%), while *Phaeocystis globosa* (4.7%) and the heterotrophic dinoflagellate *Gyrodinium fusiforme* (1.8%) were most abundant with SR (Figure 1H).

**Figure 1.**
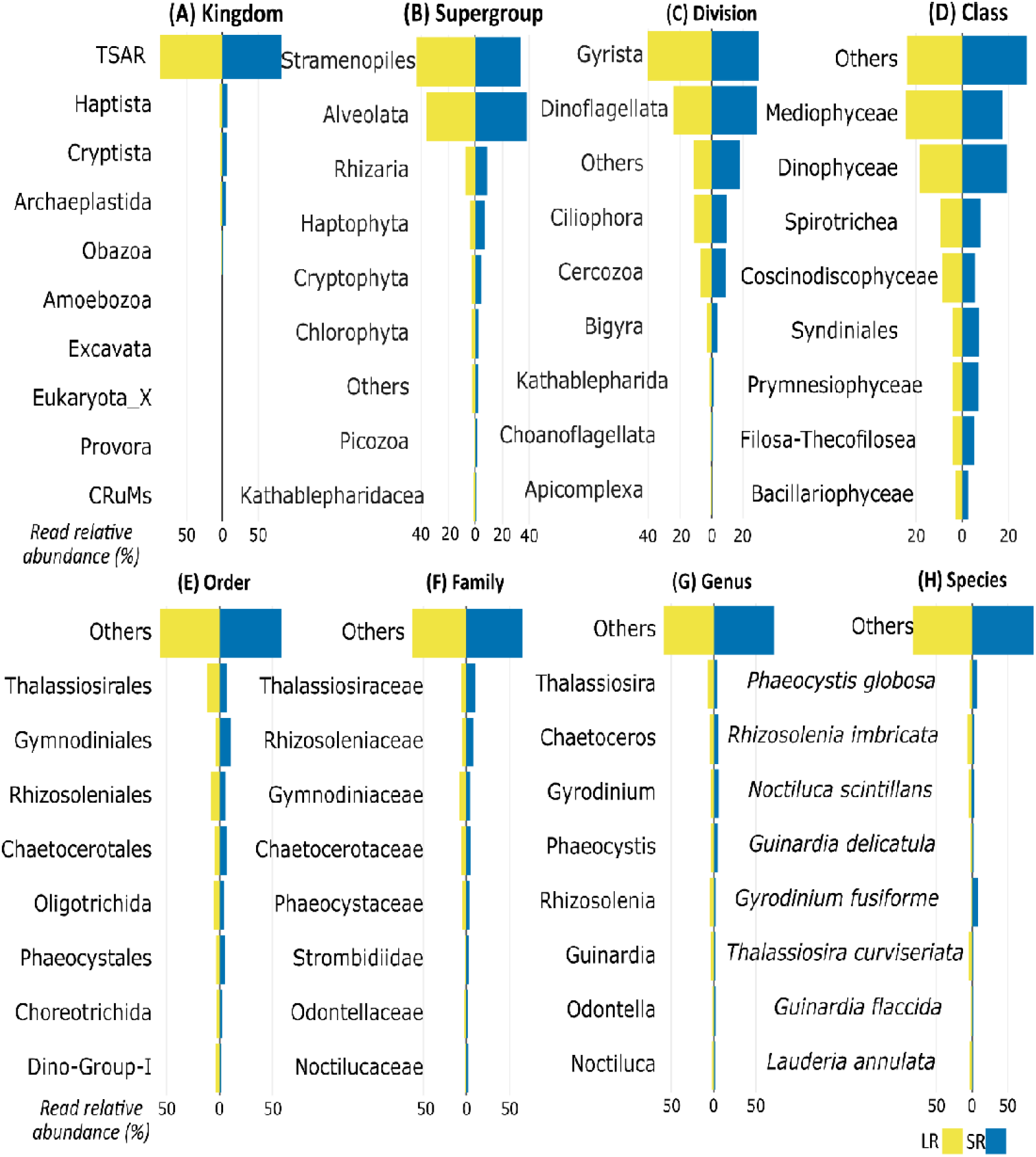
Dual-horizontal histograms of protist compositions based on read relative abundance (%) obtained through LR (yellow) and SR metabarcoding (blue) across the following taxonomic ranks: (A) Kingdom, (B) Supergroup, (C) Division, (D) Class, (E) Order, (F) Family, (G) Genus, and (H) Species, in 79 samples. In total, 7,438,382 reads were identified as protists and classified as 1,039 species based on the SR metabarcoding dataset, while 7,623,617 reads were identified as protists in 1,180 species from the LR metabarcoding dataset.

**Figure 2.**
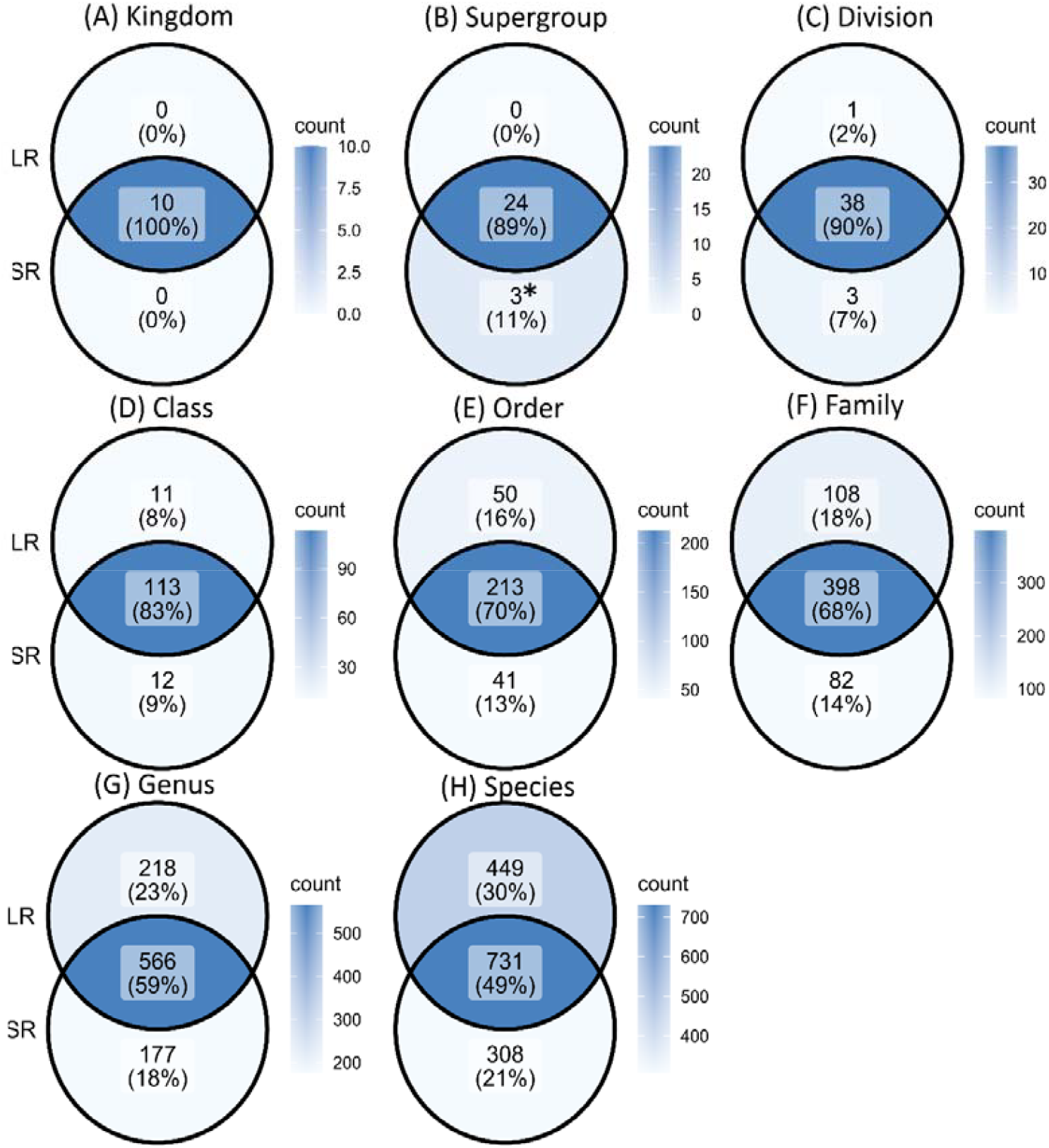
Venn diagrams illustrating the absolute number of protist taxa and their relative abundance (%) as detected with LR, and SR metabarcoding approaches across the following taxonomic ranks: (A) Kingdom, (B) Supergroup (the asterisk indicate the unique Supergroup detected in SR: Amoebozoa_X, Mantamonadidae, and Rhodelphidia), (C) Division), (D) Class, (E) Order, (F) Family, (G) Genus, and (H) Species in 79 samples. The intersecting area indicates the common taxa identified by both methods. In total, 7,438,382 reads were identified as protists and classified as 1,039 species based on the SR metabarcoding dataset, while 7,623,617 reads were identified as protists in 1,180 species from the LR metabarcoding dataset.

### Comparing the identification of unique taxa between SR and LR metabarcoding

Venn diagrams showed a high congruence between the two metabarcoding approaches in the detection of common taxa at higher taxonomic ranks, with progressively fewer shared and more unique taxa at lower taxonomic levels (Figure 2A–H). Both SR and LR datasets identified the same kingdoms (Figures 1A and 2A), and at the supergroup and division levels, approximately 90% of detected taxa were shared between the two approaches (Figures 2B–C). SR detected three unique supergroups (Figure 2B) (Amoebozoa_X, Mantamonadidae, and Rhodelphidia). The difference between the two approaches became more pronounced at the order and family levels, where 108 families (18%) were only identified in the LR dataset, whereas 82 families (14%) were unique to the SR dataset (Figure 2E–F). At genus and species levels, the percentage of common taxa further declined, with less than 60% and 50% of taxa being shared between the two methods (Figure 2G–H). Overall, the LR dataset showed a higher proportion of unique taxa from the family to species levels (Figure 2). The intersection part of the Venn diagram, illustrating the shared taxa between the two datasets, was dominated by “abundant” taxa with a relative read abundance of more than 0.1%. (Tables S2, Figure 2). In contrast, uniquely detected taxa were rare or “intermediate rare” taxa, (Table S2). Only a few unique genera were abundant (see Table S3). Eleven abundant genera were detected exclusively with LR, while three were unique to SR (Table S3). For example, only LR detected the autotrophic protists *Gephyrocapsa* (ex. *Emiliania*, Haptophyta) and *Bellerochea* (Bacillariophyceae), while the flagellates *Torodinium* and Telonemia-Group-2 were only identified with SR (Table S3). When comparing the distribution of identified rare taxa at various taxonomic levels between the two datasets, the SR dataset consisted mostly of abundant taxa, while the LR dataset contained more rare taxa (Table S2).

### Protist diversity and community structure across a coastal-offshore gradient

The diversity indexes Richness, Shannon and Gini-Simpson (1-D) showed a slightly decreasing trend from the coastal to the offshore station for both datasets, with no significant differences among locations with SR (Figure 3A–C; Tables S4, S5, S6, S7). In contrast, in the LR dataset, significant differences among locations were observed for all diversity indexes (i.e., Richness, Shannon, Gini-Simpson, Chao1; Tables S6 and S7). In contrast to Shannon and Simpson indexes, the Chao index reached higher mean values with LR (Figure 4D). Protist communities displayed the highest observed richness in the LR dataset (e.g., 455) at the coastal station MOW1 (Figure 3A, Table S6). The rarefaction curves showed that a lower sampling size is sufficient for LR to capture the same protist diversity as with SR (Figure 3E). A comparison between the mean ranks with the Wilcoxon test revealed significant differences between the two metabarcoding approaches in Shannon, Gini-Simpson and Chao1 alpha diversity indexes but not in richness (Table S7), although mean richness was slightly lower with LR (Table S6). In addition, alpha diversity indexes (except Chao1) of the LR dataset showed higher variability than those of the SR dataset (Figure 4A-E).

**Figure 3.**
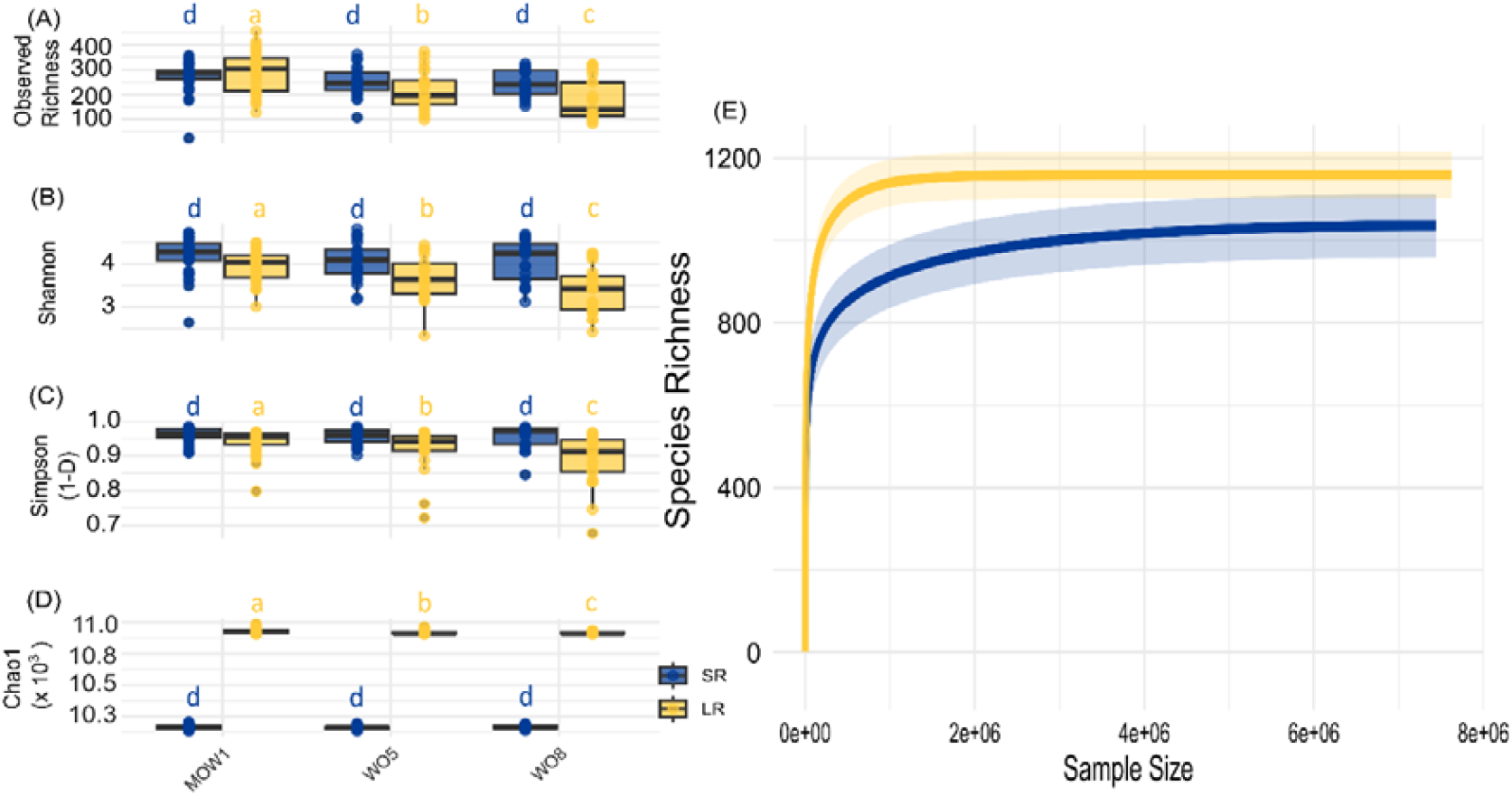
Comparison of protist species -diversity indexes in: (A) observed species richness, (B) Shannon index, (C) Gini-Simpson (1-D), (D) Chao1 index; for short-read (SR, blue) and long-read metabarcoding approaches (LR, yellow) and three different sampling locations (MOW1; WO5; WO8), (E) Rarefaction curves of protist species richness obtained through short-read (SR, blue) and long-read metabarcoding (LR, yellow) approaches with increasing read counts (i.e., sequencing depth). Significant differences between geographic locations (with Kruskall-Wallis tests) are indicated by different small letters (a, b, c), the absence of significance is indicated with the small letter d. The SR and LR datasets were rarefied at 10,131 reads per sample. Each dataset contained a total of 769,956 protist rarified reads in 76 samples. A total of 1,039 and 1,180 species were detected in SR and LR respectively.

**Figure 4.**
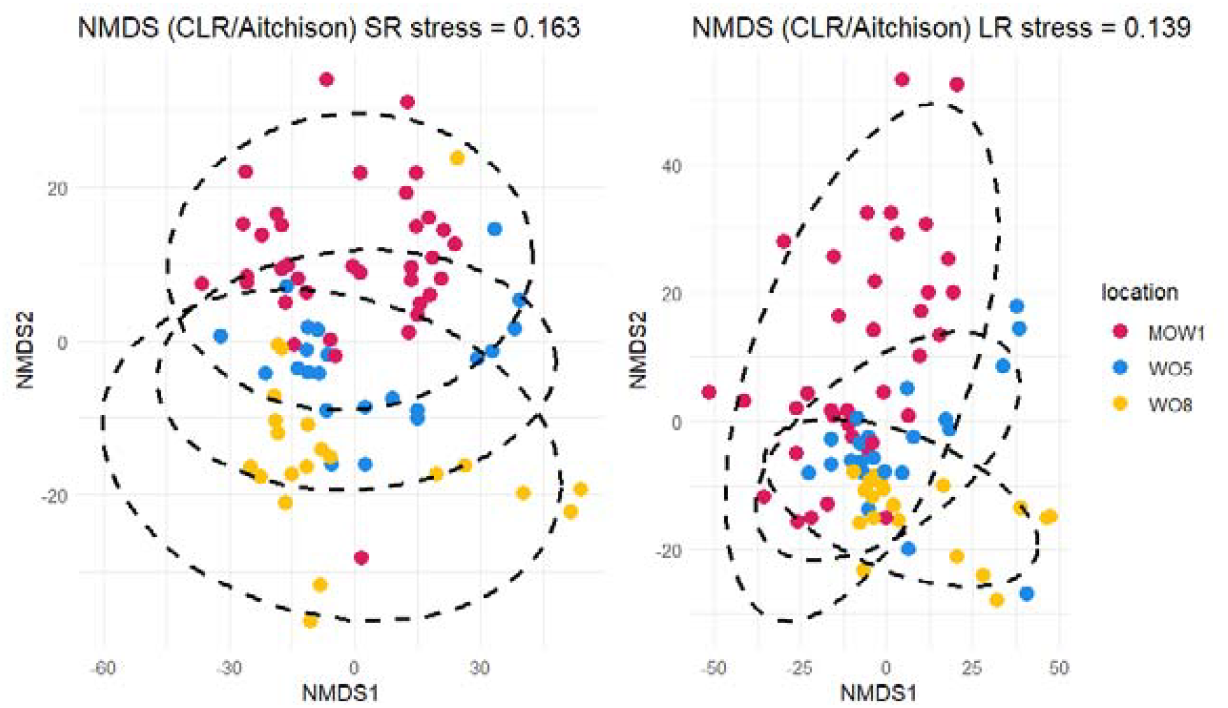
Ordination with multidimensional scaling (NMDS) and CLR transformation of protist communities for short-read (SR, left) and long-read metabarcoding (LR, right) methods and across geographic sampling locations (MOW1, pink; WO5, blue; WO8, yellow, 79 samples in total). Each dot represents one individual sample, and each colour represents a sampling station. In total, 7,438,382 reads were identified as protists and classified as 1,039 species based on the SR metabarcoding dataset, while 7,623,617 reads were identified as protists in 1,180 species from the LR metabarcoding dataset.

NMDS and PERMANOVA analyses indicated that protist communities differed significantly among stations (p < 0.001 for both datasets); however, only 7% and 9% of the total variation was explained by location in SR and LR, respectively (Figure 4). The coastal-offshore gradient was somewhat more noticeable in the NMDS of the LR dataset. When comparing spatial patterns of the composition of protist communities at the order level, both methods detected the same orders along the coastal-offshore gradient, albeit in different proportions (Figure 5). Both methods showed that the diatom order Mediophyceae dominated the coastal station while at the transitional and offshore stations, Dinophyceae were more abundant, again with both SR and LR.

**Figure 5.**
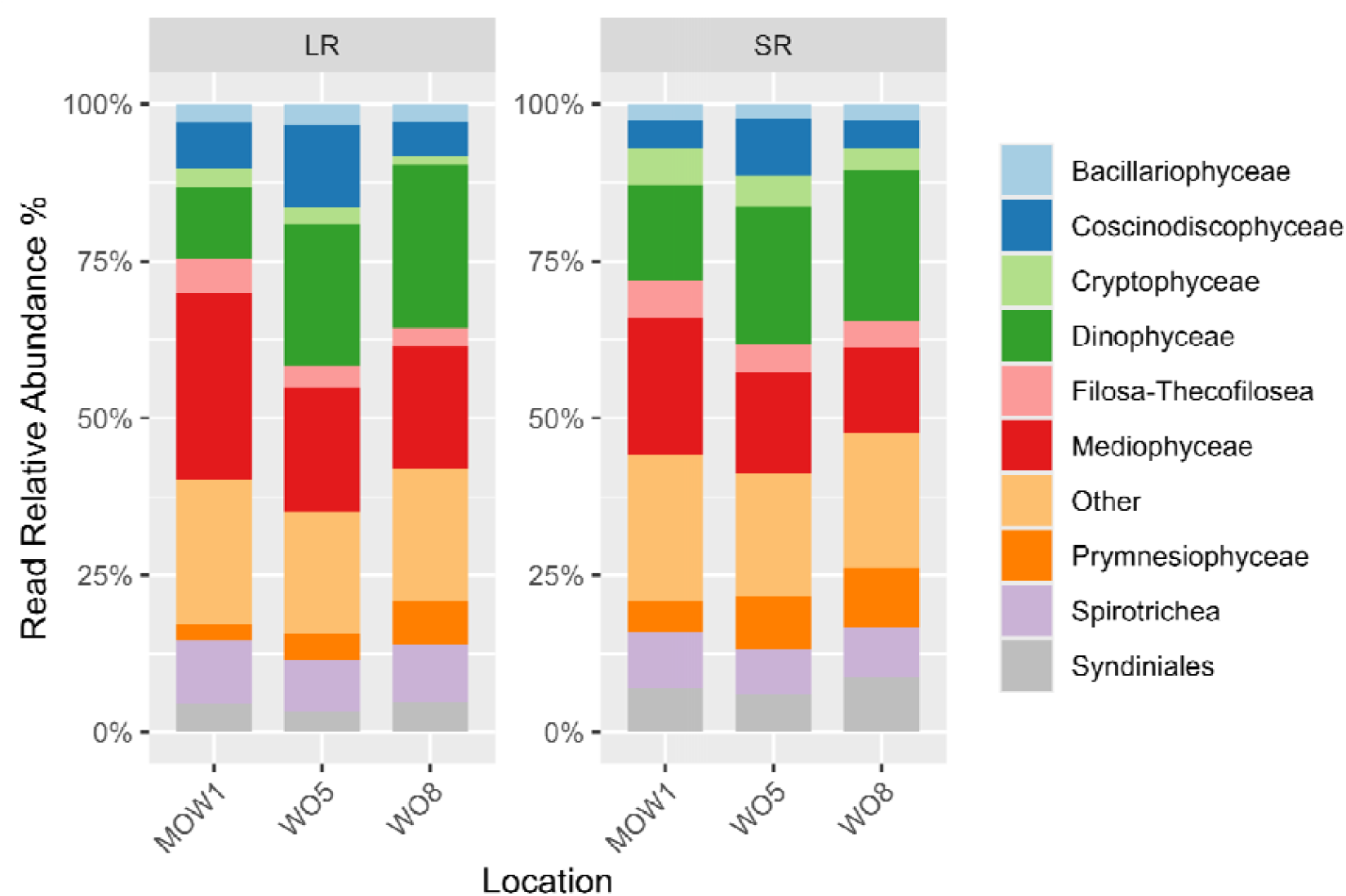
Relative read abundance of protist reads for short-read (SR, right) and long-read metabarcoding (LR, left) techniques and the three sampling locations (MOW1; WO5; WO8) in 79 samples. In total, 7,438,382 reads were identified as protists and classified as 1,039 species based on the SR metabarcoding dataset, while 7,623,617 reads were identified as protists in 1,180 species from the LR metabarcoding dataset.

## Discussion

### Sequencing yield

Regarding overall sequencing yields, both SR and LR metabarcoding approaches were equally suitable, with LR producing somewhat higher overall numbers of sequencing reads than SR (22 million vs 17 million raw reads, and 14 million as compared to 13 million filtered reads). If only sequencing reads being assigned to protists (∼7.6 million vs ∼7.4 million) and their species richness (1,180 vs 1,039) were considered, again, both approaches performed alike and showed a dominance of protists. A slightly greater relative abundance of protists (>10%) was observed with SR than with LR (Figure S12).

### Taxonomic assignment

A higher proportion of unassigned taxa was observed with the SR method (Figure S13), matching the metabarcoding results of protist communities along the coast of Texas (Gaonkar and Campbell, 2024). Furthermore, taxa identified by both methods predominately belonged to higher taxonomic levels (Figure 2), which is consistent with findings by Gaonkar and Campbell (2024). In the latter study, sequencing reads covering the entire 18S rDNA region provided more accurate taxonomic assignments with fewer unassigned reads when compared to results based on the V4 (380bp) and V8-V9 regions (330bp). Although our study amplified only a part of the 18S rDNA region using both methods, LR still provided more reliable taxonomic identifications than SR (i.e., SR: 15.2 % vs LR: 1.2 % unassigned species). This suggests that the longer 18S rDNA fragment being analysed by the LR approach in our study was more effective in capturing a broad range of taxa. Differences in taxonomic assignment between short-read (SR) and long-read (LR) datasets may partly be attributed to different bioinformatic pipelines including different classifier methods (Mugnai et al. 2023; Hleap et al. 2021). Expectation–maximization algorithms such as EMU can distinguish closely related taxa or assign unknown sequences to their nearest available reference (Curry et al., 2022). This probabilistic framework is designed to account for sequencing errors inherent to long-read sequencing data. Unlike the naïve Bayesian classifier used for SR data with DADA2, EMU does not apply a fixed bootstrap confidence cutoff but instead estimates relative abundances based on probabilistic alignment to reference sequences. Consequently, EMU may yield deeper taxonomic assignments with the same reference database, even when sequence identity is lower. As EMU tends to assign reads to the nearest available reference even with limited similarity, whereas the naïve Bayesian classifier that was used for SR analyses truncates assignments when confidence is low, a higher proportion of species-level assignments in LR data may reflect algorithmic permissiveness. In our study, the use of different classifiers followed commonly applied, technology-specific workflows for SR and LR metabarcoding.

In addition, it is worth noting that while recent improvements in Oxford Nanopore library kits and flow cell chemistry have substantially reduced chimeras and sequencing error rates (Wang et al., 2021), PCR chimeras, and systematic indel and homopolymer errors might remain potential problems in LR amplicon datasets and may contribute to inflated richness or spurious species-level assignments. The EMU pipeline used in this study is optimized for error-prone reads, yet it does not include an explicit chimera detection or removal step, Consequently, undetected chimeras may have contributed to elevated richness or deeper taxonomic assignments in the LR dataset, and results at the species level should therefore be interpreted with caution. Reference based chimera detection tools such ‘uchime2_ref’ have been applied to higher quality long reads produced by PacBio or Nanopore using unique molecular identifiers during library preparation before sequencing (Karst et al., 2021). However, these approaches might remove genuine biological sequences or fail to detect chimeras when sequencing error rates are high (Stock et al., 2025). Finally, custom pipelines such as CONCOMPRA have been recently applied to the analyses of prokaryotic communities, but to our knowledge these approaches have not yet been implemented in eukaryotic long-read metabarcoding (Stock et al., 2025).

Differences in the detection of protist taxa between the two methods were particularly evident at finer (lower) taxonomic levels (Figure 2). Both datasets recovered most of the abundant taxa, while unshared taxa were typically rare (Table S3). Notably, more rare taxa were detected in the LR metabarcoding dataset (Table S2). This is consistent with previous findings that ONT long sequencing was better suited for detecting rare taxa in a microbiome study (Szoboszlay *et al*., 2023). To account for the potentially higher detection of rare taxa in our study, we applied an abundance threshold. Without this control, the probabilistic model used by the EMU algorithm in LR analyses might artificially inflate the number of rare species (Curry *et al*., 2022).

### Composition differences and identification of specific taxa

We expected congruence between the SR and LR metabarcoding approaches in the relative read abundance at high taxonomic levels (Figure 1), in line with the findings by Gaonkar and Campbell (2024). However, at lower taxonomic levels, notable differences in protist relative abundance were observed between the two methods. Specifically, Dinophyceae (dinoflagellates), and closely related groups (e.g., Syndiniales), were more abundant in the SR dataset, while the LR dataset showed a prevalence of Bacillariophyceae (diatoms, Figure 1). Our results thus confirm one of the known limitations of SR metabarcoding; the over-presentation of dinoflagellates due to their high rDNA copy numbers (Prokopowich *et al*., 2003; Georges *et al*., 2014; Bradley *et al*. 2016 and reference therein; Santi *et al*., 2021; Yeh *et al*., 2021). This limitation is especially relevant in diatom-dominated marine systems such as the BPNS (Nohe *et al*., 2020). In contrast, the LR approach appeared to mitigate this bias, probably due to several possible reasons, besides the differences in bioinformatic pipelines mentioned in the previous section: (1) Technological differences: SR methods rely on bridge amplification in Illumina sequencing, which might introduce amplification bias. In contrast, LR sequencing uses ligation and direct sequencing, which may reduce such bias (Mikheyev and Tin, 2014) and help correct the over-representation of dinoflagellates. (2) Completeness of reference database: The dinoflagellate reference database remains limited at the genus/species level, potentially further contributing to biased classifications with SR methods (Gaonkar and Campbell, 2024, Mordret *et al*., 2023). Conversely, diatom reference databases are more complete, enhancing the LR’s ability to accurately detect and classify them (Gaonkar and Campbell, 2024). An alternative approach to address database limitations would be taxonomy-free assignments, such as de novo assembly. While promising and already applied to microbial profiling (Stock *et al*., 2025) these approaches currently require metagenomic or long read sequencing data. (3) Intracellular haplotype diversity: Certain dinoflagellates, such as *Tripos sp*., exhibit high single-cell haplotype diversity, where a single cell may contain multiple haplotypes (Huang *et al*., 2024). In our study, *Tripos* was a dominant genus (ca. 0.1 % in relative abundance) in both datasets. Such intra-cellular variation can affect the clustering of short reads, leading to inflated diversity at lower taxonomic levels by misclassifying intraspecific variation, as separate OTUs (Operational Taxonomic Units) (Stoeck *et al*. 2024; Huang *et al*., 2024). We minimized this bias by using non-clustering techniques such as amplicon sequence variants (ASVs) for SR and EMUs for LR. (4) Primer bias: Primers targeting specific parts of the 18S rDNA gene often have narrow taxonomic coverage, which can affect detection efficiency (Vaulot *et al*., 2022) and could have further contributed to discrepancies between the two methods.

Interestingly, while the SR approach identified a greater number of abundant species, the LR method detected a higher number of unique, abundant taxa (Table S2). This difference is particularly important, as only the LR approach identified key taxa that have been widely reported in the BPNS using conventional methods. Two notable autotrophic protists are the colonial diatom *Bellerochea* and the calcifying haptophyte *Gephyrocapsa* (Table S3). *Bellerochea* blooms have been recently intensified in the BPNS, likely due to temperature-driven declines in copepod populations (Mortelmans *et al*., 2024). The future intensification of these blooms is alarming as they might reduce oxygen levels in the water column, potentially threatening higher trophic levels, including larval fish (Mortelmans *et al*., 2024). *Gephyrocapsa*, on the other hand, is ecologically important due to its substantial production of biomass and calcium carbonate. It contributes to carbon dioxide uptake, and emits dimethyl sulphide, a gas with climate-cooling properties (Paasche, 2001). This haptophyte forms extensive blooms that can span thousands of kilometres in summer and autumn (Raitsos *et al*., 2006). These blooms can occur locally in the North Sea (Holligan *et al*., 1993), or they can be further transported via the inflow of Atlantic waters (Head *et al*., 1998). To date, most research on this haptophyte has relied on satellite data to characterize its spatial distribution (e.g., Ladd *et al*., 2018; Terrats *et al*., 2020), while studies on its phenology using eDNA approaches remain very limited (Neri *et al*., 2025). Future studies on *Gephyrocapsa* in the BPNS should integrate eDNA and satellite technologies to better resolve the spatial and temporal dynamics of this haptophyte at local and regional scales.

### Coastal-offshore gradient influencing protist diversity and community structure

Although both SR and LR methods revealed similar trends in protist diversity patterns, community structure, and composition at higher taxonomic levels along the coastal-offshore gradient (Figures 3, 4, 5, and Table S1), notable differences emerged in the estimated diversity metrics between the two methods (Figures 3 and 5). Given the increased sequence context provided by LR metabarcoding, we initially expected LR to yield deeper taxonomic annotation and potentially identify a higher number of taxa under the applied workflow compared to SR. Discrepancies in diversity estimates can at least partially be explained by the higher number of rare taxa identified by LR, as mentioned in the previous section.

Because the Shannon diversity index is sensitive to the presence of rare taxa (Roswell *et al*., 2021), the prominence of such rare taxa in the LR dataset most likely influenced this index (Figure 3B; Table S2). Similarly, the Gini-Simpson index, which emphasizes taxon dominance, was lower in the LR dataset, reflecting its detection of fewer dominant taxa compared to SR (Figure 3C; Table S2; Roswell *et al*., 2021). In contrast, the Chao1 index (Figure 3D), which estimates species richness while correcting for rare taxa, may provide a more reliable estimate of protist richness in the LR dataset (Chao *et al*., 2006; 2020). Chao1’s mathematically framework reduces bias resulting from underrepresentation of rare species, making it particularly suited for use with high-resolution sequencing data (Figure 3D; Table S8).

Similar inconsistencies in diversity estimates between SR and LR methods have been reported in microbiome research. For instance, Szoboszlay *et al*. (2023) found that SR data tended to overestimate bacterial diversity in faecal samples, likely due to sequencing noise. These findings emphasize the importance of selecting appropriate diversity indexes based both on the metabarcoding method used and the nature of the data.

Phylogenetic-based metrics such as the Faith’s Phylogenetic diversity index (PD; Faith, 1992) and UniFrac (Lozupone and Knight, 2005) may offer more accurate insights for metabarcoding data by incorporating evolutionary relationships. However, the primary aim of the current study was not to evaluate all available diversity indexes, but rather to provide a first comparison of the most commonly used indexes within the context of eDNA-based protist community analyses.

In contrast to the diversity indexes, protist community structure differed significantly across sampling stations, although geographic location accounted for less than 10 % of the variability for both methods (Figure 4). Nonetheless, the differences in protist communities along the coastal-offshore gradient were more pronounced in the LR dataset. This pattern may be attributed to the greater depth of taxonomic assignment and the detection of additional low-abundance taxa under the LR metabarcoding workflow, which likely captured subtle shifts in community composition. At higher taxonomic level, both methods provided consistently detected a dominance of Mediophyceae (diatoms) at the coastal station, while Dinophyceae (dinoflagellates) prevailed at the transient and offshore stations (Figure 5). The dominance of diatoms at the coastal station MOW1 can probably be explained by riverine inputs from the River Schelde (Aubert *et al*., 2022).

### Recommendations

Given the inherent limitations of each method, the choice between SR and LR metabarcoding for marine protist monitoring studies depends heavily on the goals of the study. SR metabarcoding remains a reliable and cost-effective tool for detecting abundant taxa and providing broad snapshots of community diversity. Its use in monitoring for almost a decade (e.g., Teeling *et al*., 2016; Lambert *et al*., 2019; Caracciolo *et al*., 2022; Skouroliakou *et al*., 2024) has enable consistent data generation over time. One of its key advantages is its established use.

As highlighted by Ducklow *et al*. (2009), long-term oceanographic surveys are crucial for capturing episodic events and assessing the impacts of climate change, processes that often unfold over extended timescales. Recent initiatives, such as the European Marine Omics Biodiversity Observation Network (EMO BON) represent promising initiatives to standardize and sustain SR-based eDNA monitoring across European waters (Santi *et al*., 2023).

Despite its advantages, SR metabarcoding has well-known limitations, including amplification bias, limited depth of taxonomic assignment, and the overrepresentation of dinoflagellates due to high rDNA copy numbers. In contrast, LR metabarcoding (particularly using Oxford Nanopore Technologies (ONT)) is emerging as a powerful complement to SR approaches. We would recommend additional verification of species assignments with LR approaches in future studies with other analyses like BLAST searches, alignments or phylogenetics, and suggest to also use mock communities or positive controls but these are beyond the scope of the present study. Recent advances in ONT accuracy and throughput are addressing previous limitations related to error rates, making LR sequencing more accessible and suitable for marine studies (Wang et al., 2021). The application of long-read (LR) metabarcoding in monitoring initiatives is promising and offers several distinct advantages: Unlike SR, LR metabarcoding might provide increased sequence context and, under certain workflows, offer deeper taxonomic assignment that may facilitate differentiation among closely related taxa; it may reduce amplification bias, resulting in more balanced community profiles; and it can enhance the detection of low-abundance taxa, thereby increasing sensitivity to subtle shifts in biodiversity. Last but not least, LR techniques quantify key phytoplankton genera more accurately, including diatoms and haptophytes such *Bellerochea* and *Gephyrocapsa*, with pronounced harmful effects in the BPNS and other parts of the North Sea.

### Conclusions

Molecular methods for the long-term monitoring of marine ecosystems continue to evolve alongside advances in sequencing technologies. Although still underutilized in protist research, long-read sequencing (LR) offers significant potential to address limitations of traditional short-read (SR) metabarcoding. In the current study, we directly compared protist diversity, taxa detection, composition, and community structure using LR and SR metabarcoding approaches. While both metabarcoding methods showed consistent patterns in community composition at higher taxonomic ranks and similar trends along the coastal-offshore gradient in the BPNS, data from LR revealed less overrepresentation of dinoflagellates and a greater capacity to detect certain diatoms with harmful effects and haptophytes in general. These results underscore the value of integrating LR sequencing into future marine eDNA monitoring frameworks, particularly for tracking harmful and/or nuisance taxa and assessing fine-scale biodiversity dynamics across spatial gradients.

## Acknowledgments

We acknowledge funding from the European Maritime, Fisheries and Aquaculture Fund (EMFAF) for the ZeroImpact project. We also thank Sophie Derycke for coordinating ZeroImpact and the Freshwater Biology team at the Royal Belgian Institute of Natural Sciences (RBINS) for their involvement in various sampling campaigns. This research was supported by the Belgian Science Policy (BELSO) within the BRAIN-be program (BG-PART, contract number *B2/202/P1/BG-PART*), PiNS, contract number *RV/21/PiNS*, and EMBRC Belgium - FWO international research infrastructure *I001621N* and *I000825N* (infrastructural funding for PAE). We thank Marie Cours, Jeroen Venderickx, the crew of RV Belgica, and all other participants in the field campaigns for their valuable technical support. We acknowledge Willem Stock for his useful insights on bioinformatic analyses.

## Data accessibility and Benefit-Sharing

Bioinformatic workflow for SR and LR analyses, processed input tables and scripts for all figures are available to the Zenodo repository: https://doi.org/10.5281/zenodo.18366431

Raw sequencing SR and LR data have been submitted and will be publicly available upon acceptance of the manuscript the Sequence Read Archive of NCBI under BioProject number PRJNA1282445.

## Author contributions

I.S. and K.S. conceptualized, supervised and provided resources for the research project. Lab analyses and/or sampling were conducted by S.D. and D.D.W.E. Data curation was done by D-I.S., D.D.W.E, and J.V. Formal analyses and visualisations od data were done by D-I.S. and D.D.W.E. The initial draft of the manuscript was done by D-I. S. The first two authors overall contributed equally. All co-authors provided detailed revisions of the manuscript. All authors contributed to or commented on the writing of the manuscript; they all have read and agreed to the submitted version of the manuscript.

## Notes

https://doi.org/10.5281/zenodo.18366431

### Competing Interest Statement

The authors have declared no competing interest.

