## Supplementary Figures for "Unveiling protist composition and diversity patterns with eDNA metabarcoding: comparing short- and long-read approaches"

***1*** *Ifremer, DYNECO, 29280 Plouzané, France*

***2*** *Laboratory of Protistology and Aquatic Ecology, Department of Biology, Ghent University, Krijgslaan 281-S8, 9000 Gent, Belgium*

***3*** *Operational Directorate Natural Environment, Aquatic and Terrestrial Ecology, Freshwater Biology, Royal Belgian Institute of Natural Sciences, Vautierstraat 29, 1000 Brussels, Belgium*

***4*** *Laboratory of Marine Biology, CP160/15 Université Libre de Bruxelles (ULB)*

***5*** *Research Group Zoology, University of Hasselt, Ago*

*ralaan Building D, 3590 Diepenbeek, Belgium*

**The first two authors contributed equally*

**Supplementary information**

**Supplementary Figures**

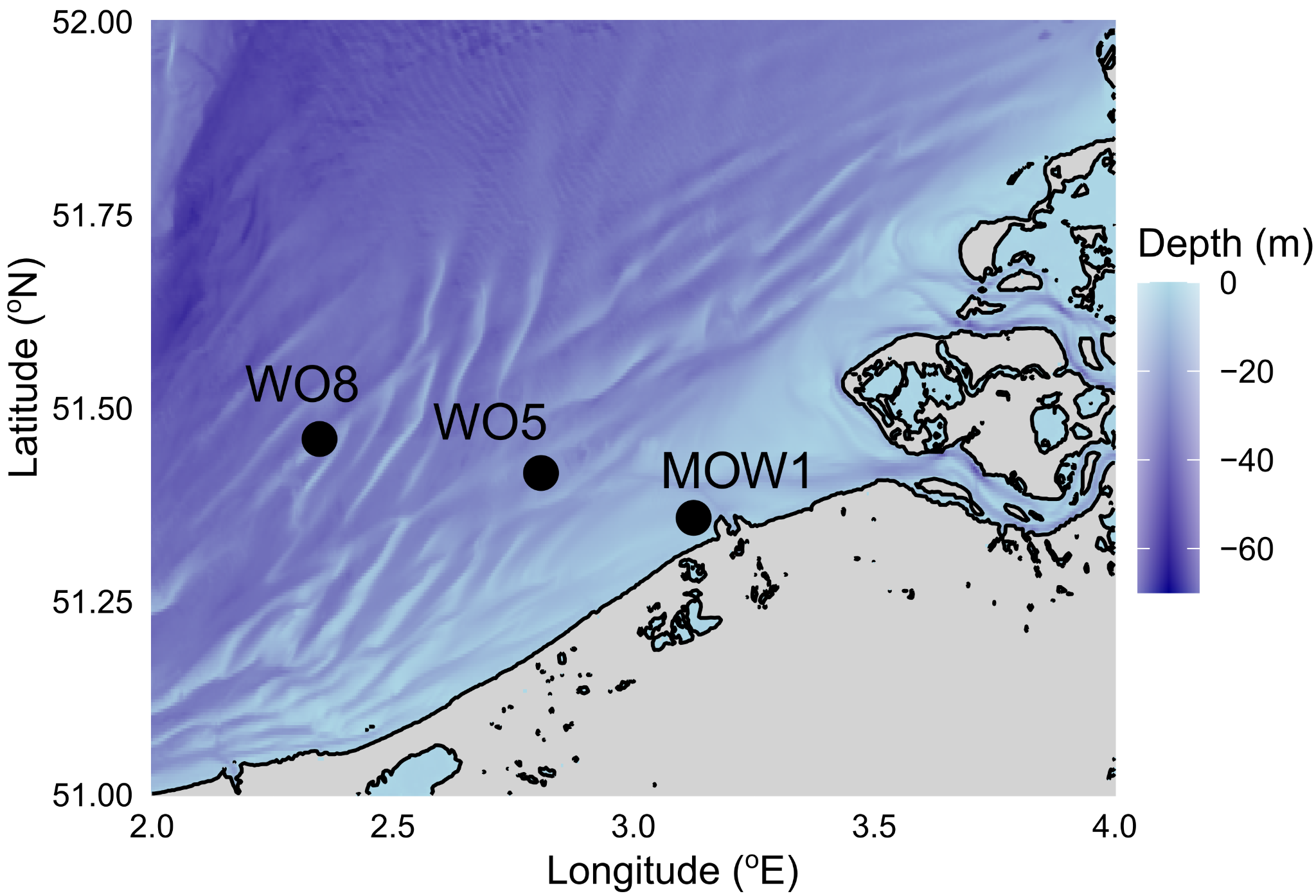

Figure S1. Bathymetry map of the Belgian part of the North Sea and the three sampling stations along the cross-shore gradient (M0W1, W05, W08). See also Table 1 for additional information.

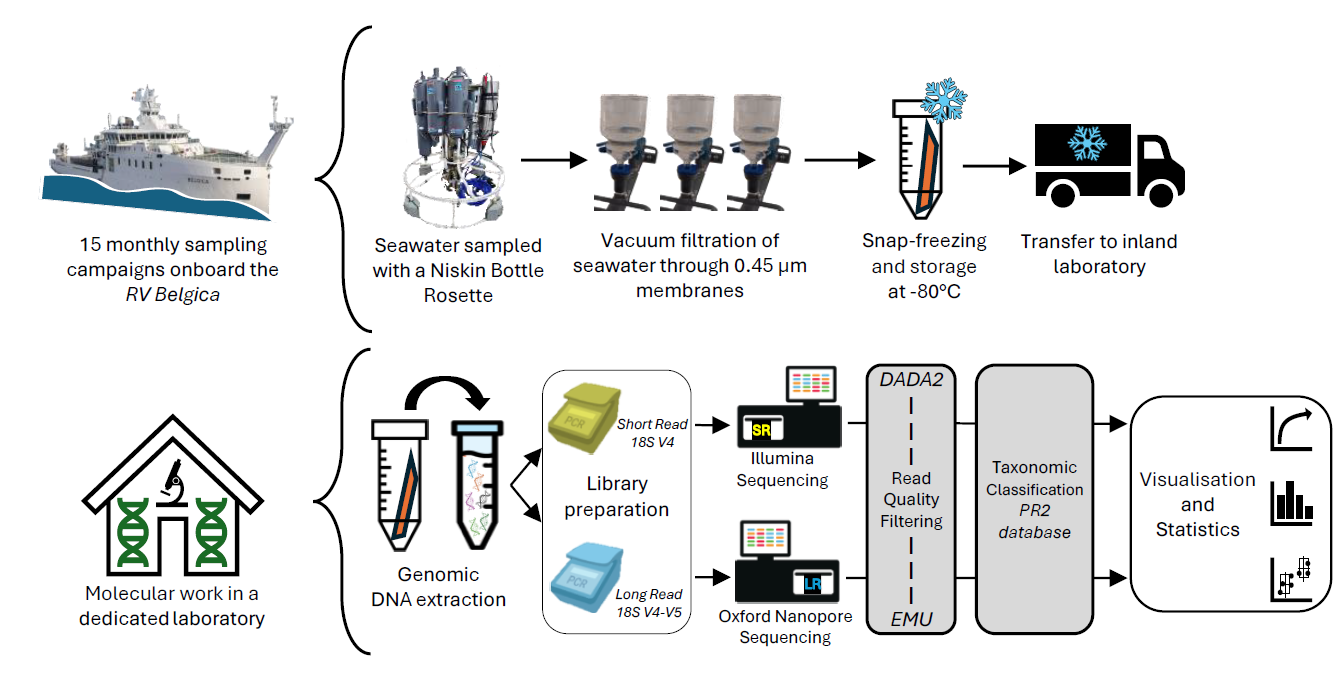

Figure S2. Overview of the sampling strategy and metabarcoding workflow: Seventy-nine samples were collected aboard the RV *Belgica* using a rosette of Niskin bottles. Seawater was filtered on board through 0.45 μm filter membranes with a low-pressure vacuum pump until saturation of the membranes. The samples were stored at −80°C on board of the vessel after being snap-frozen in liquid nitrogen and later transferred to the Royal Belgian Institute of Natural Sciences (RBINS) for DNA extraction. Libraries for long-read (LR) metabarcoding were prepared at RBINS, and short-read (SR) metabarcoding libraries were prepared at Ghent University. Sequencing was performed by OHMX (Ghent, Belgium) and Genewiz (Germany GmbH, Leipzig) for LR and SR, respectively. Analyses of sequences was conducted with DADA2 (Callahan *et al*., 2016) and EMU (Curry *et al*., 2022) for SR and LR data, respectively. The PR2 database was used for taxonomic assignment of both datasets. Statistical analyses and visualizations were performed in R (for more details, see Materials and Methods).

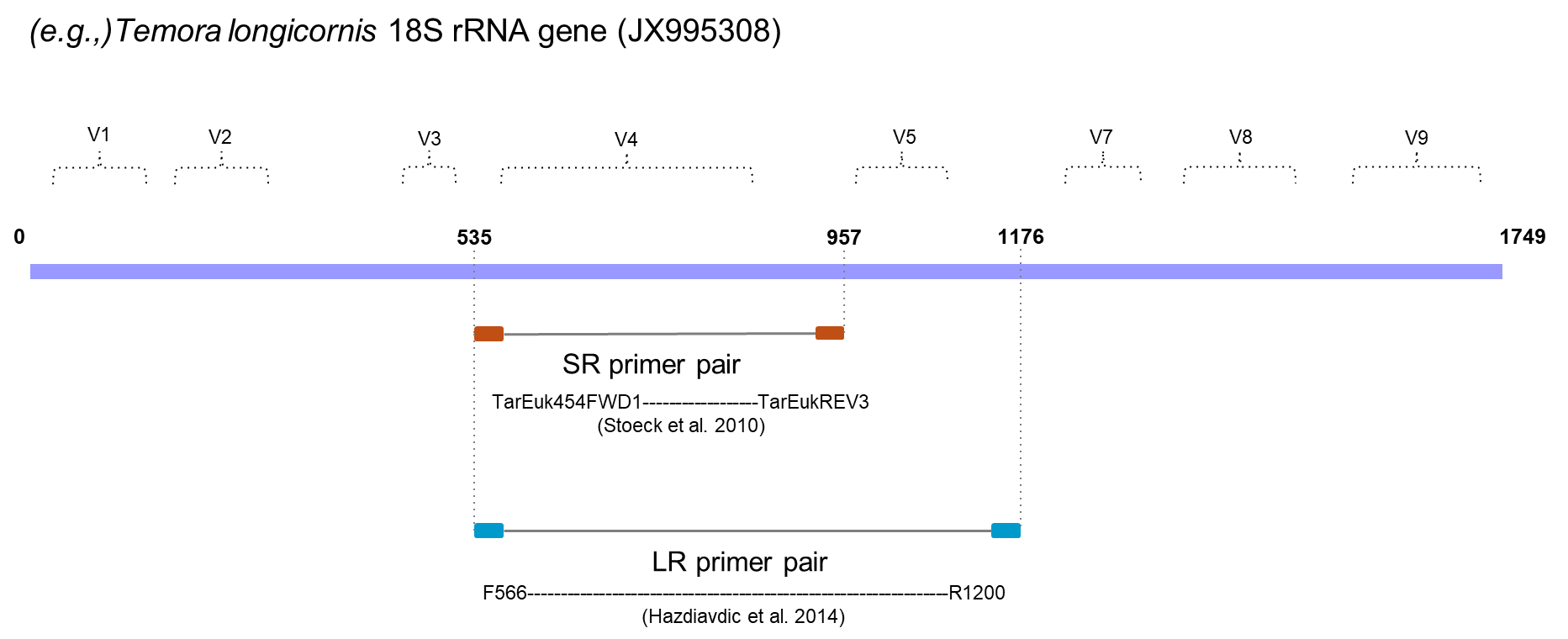
Figure S3. Schematic representation of the 18S ribosomal RNA region of the copepod *Temora longicornis* (JX995308). The position of variable regions (V1 to V9) and the overlapping primer pairs for short-read (SR; TarEuk454FWD1 – TarEukREV; generating 577 bp amplicon size) and long-read (LR; F566 – R1200; generating 745 bp amplicon size) metabarcoding (Stoeck *et al*., 2010; Hazdiavdic *et al*., 2014) are indicated along the region. Due to the low quality of reverse reads, only the forward reads of SR sequencing reads (generating 280 bp amplicon size) were kept for downstream analysis (Figs. S6A-B).

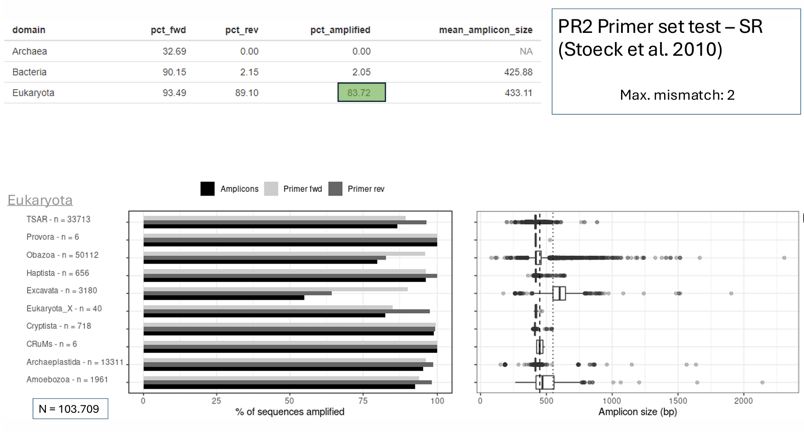
Figure S4. Summary of *in silico* tests performed on the primers TarEukFWD and TarEukREV3 (Stoeck *et al*., 2010), using the PR2 Primer test interface. **Top left**: Table summarizing the percentage of taxa of the three domains (Archea, Bacteria and Eukaryota) being amplified with either the forward (pct_fwd), or the reverse (pct_rev) primer, or the combined pair (pct_amplified). The latter notably amplified 83.72% of Eukaryota. **Bottom Left**: Horizontal bar plot displaying the percentage of sequences of various protist groups amplified with the primer pair (black), the forward (light grey) and the reverse (grey) primer. N refers to the total number of eukaryotic sequences present in the PR2 database. **Bottom right**: The *in silico* amplicon size is visualized in a box plot, with jitter dots following the legend colour and the taxonomic grouping.

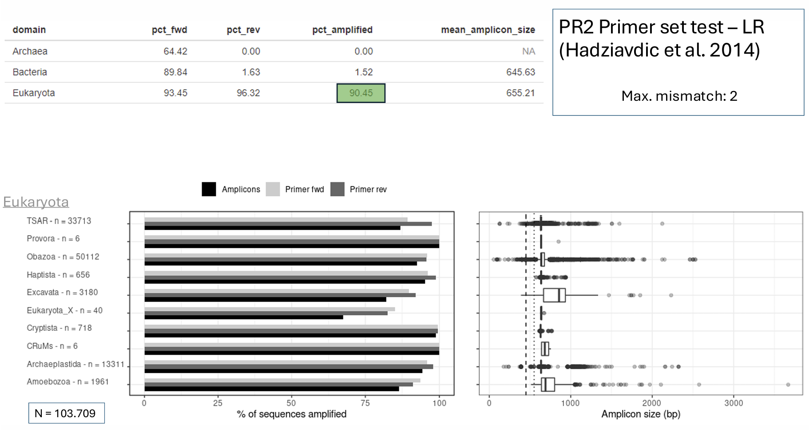

Figure S5**.** Summary of *in silico* tests performed on the primers F566 and R1200 (Hadziavdic *et al*., 2014), using the PR2 Primer test interface. **Top left**: Table summarizing the percentage of taxa of the three domains (Archea, Bacteria and Eukaryota) being amplified with either the forward (pct_fwd), or the reverse (pct_rev) primer, or the combined pair (pct_amplified). The latter notably amplified 90.45% of Eukaryota. **Bottom Left**: Horizontal bar plot displaying the percentage of sequences of various protist groups amplified with the primer pair (black), the forward (light grey) and the reverse (grey) primer. N refers to the total number of eukaryotic sequences present in the PR2 database. **Bottom right**: The *in silico* amplicon size is visualized in a box plot, with jitter dots following the legend colour and the taxonomic grouping.

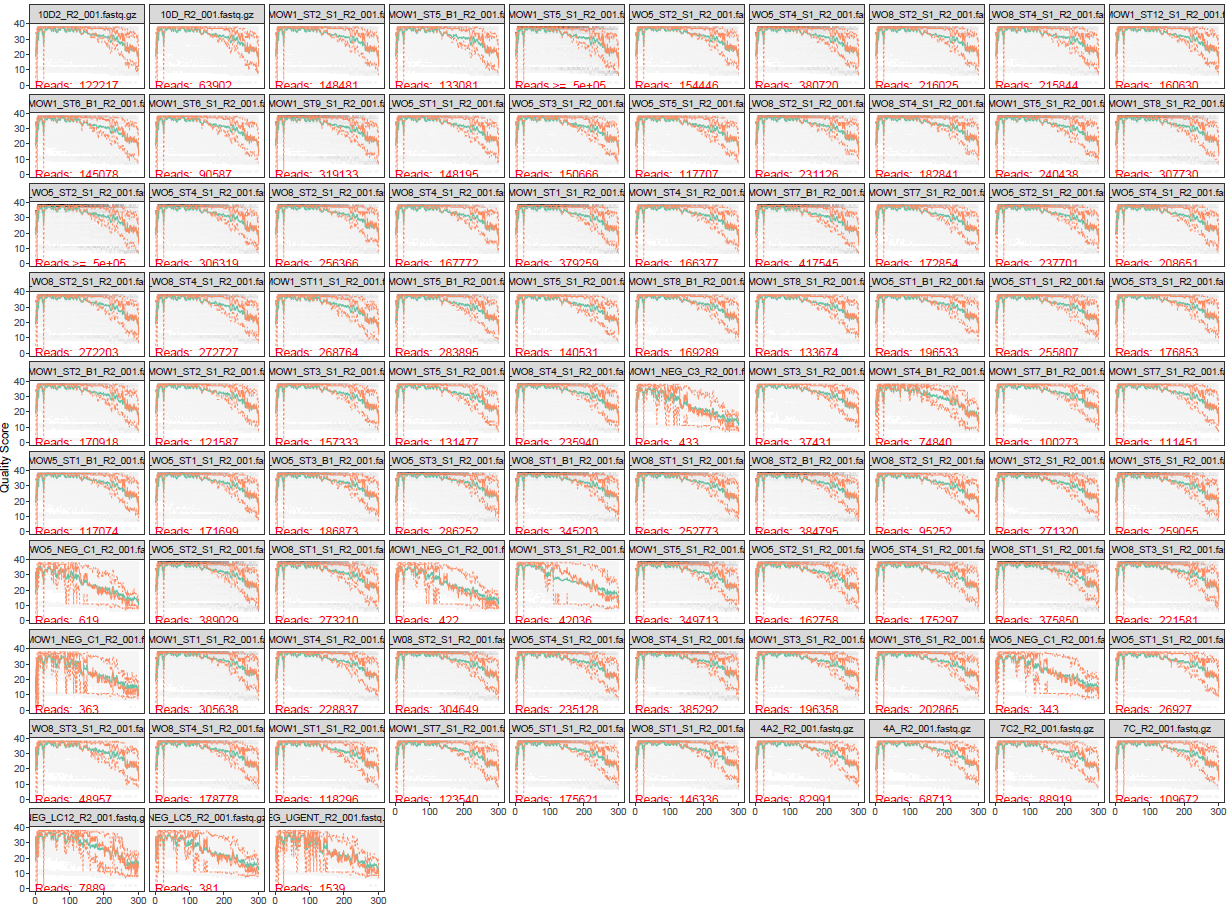

Figure S6. (A) Quality plots of the reverse sequencing reads of 79 samples and 8 negative controls. The gray scale provides a heat map of the frequency of each quality score at each base position. The mean quality score at each position is indicated by the green line, and the distribution of the quartiles of the quality score is illustrated by the orange lines.

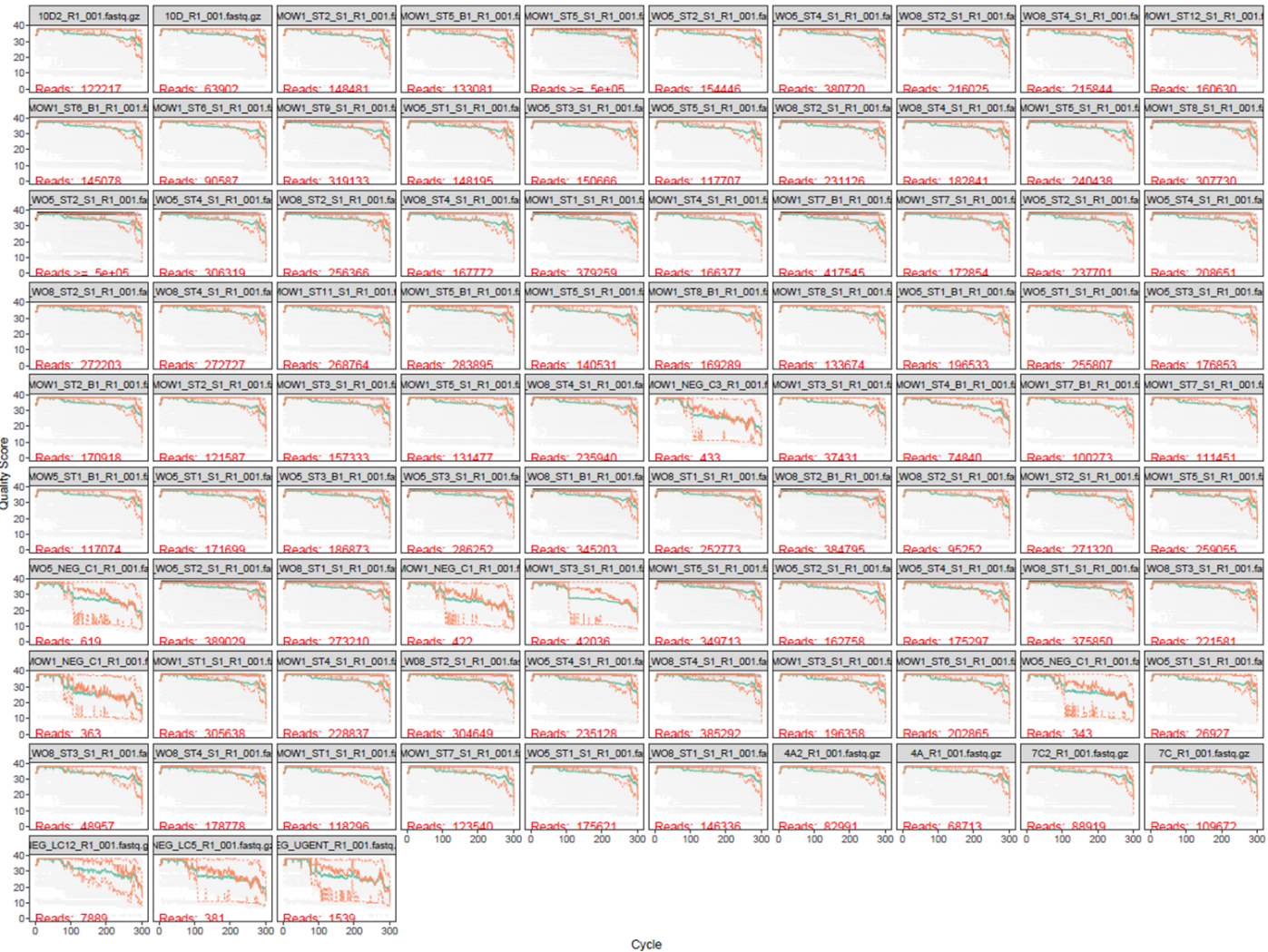

Figure S6. (B) Quality plots of the forward sequencing reads of 79 samples and 8 negative controls. The gray scale provides a heat map of the frequency of each quality score at each base position. The mean quality score at each position is indicated by the green line, and the distribution of the quartiles of the quality score is illustrated by the orange lines.

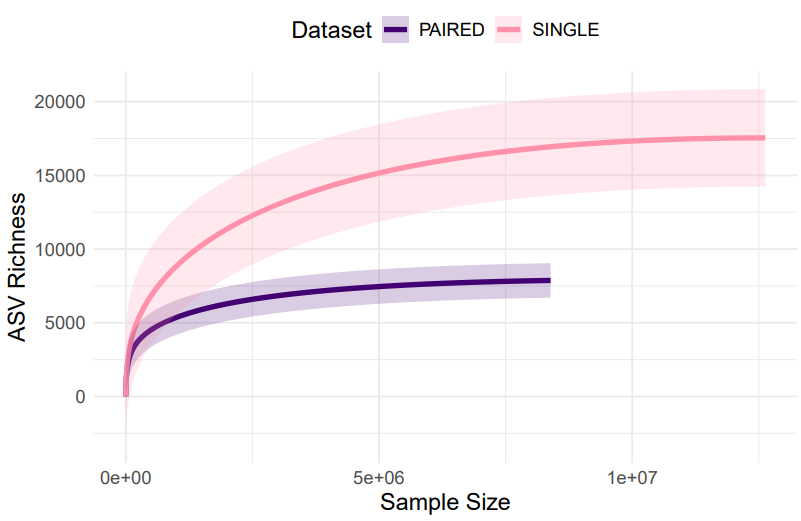

Figure S7. Rarefaction curves of Amplicon Sequence Variants (ASV) being identified as eukaryotes with SR based on both forward and reverse reads (paired-end) in purple or only forward (single-end) reads in pink.

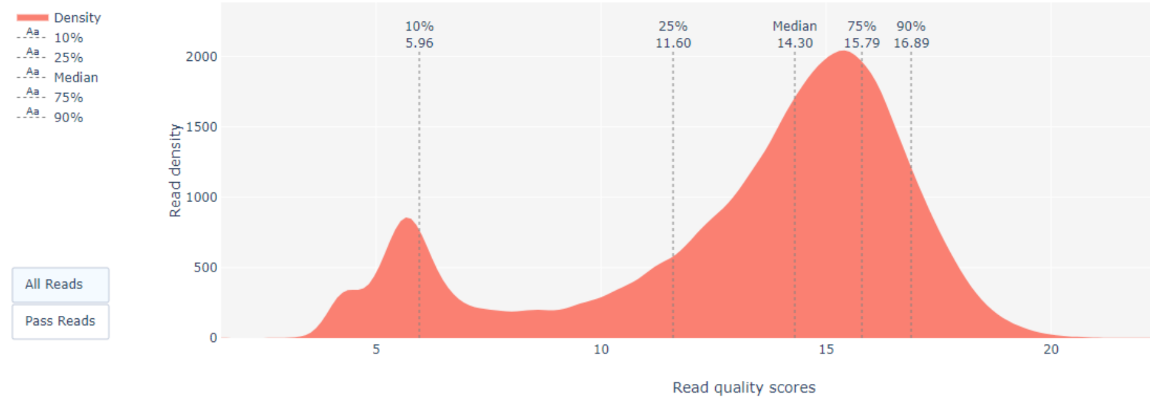

Figure S8. Basecalled PHRED read quality distribution of long read sequencing run in the entire dataset (87 samples: 79 field samples and 8 negative controls) provided by ohmx.bio dry lab report. The median quality score of the sequencing run was 14.3.

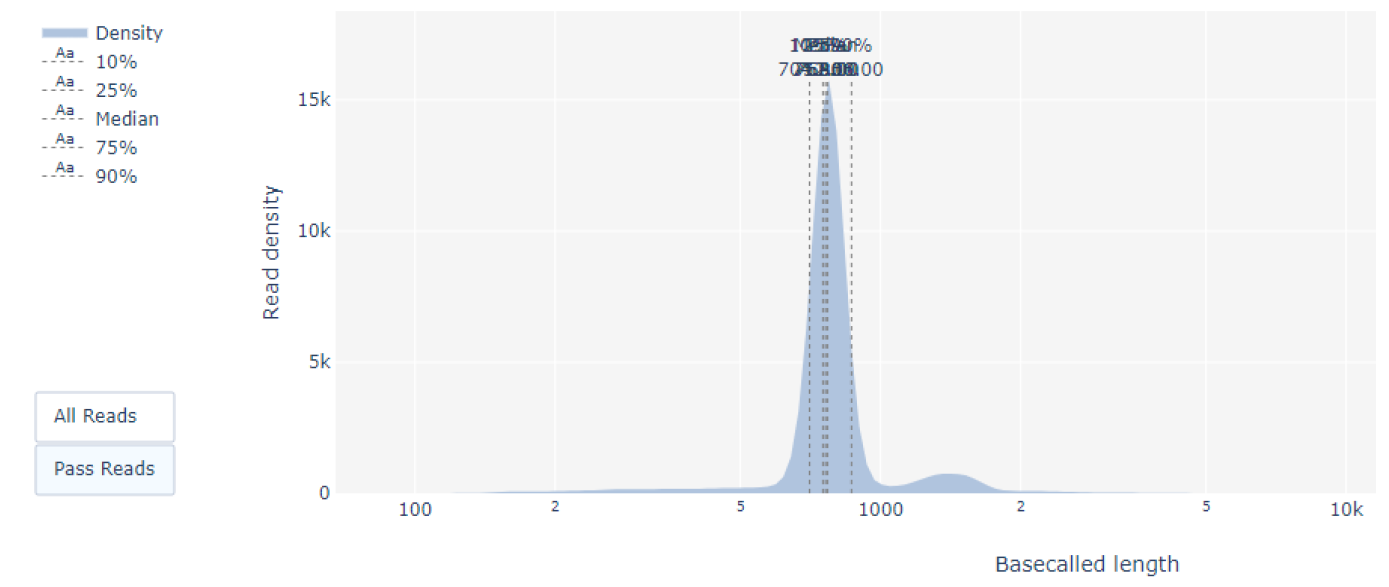

Figure S9. Long Read length distribution of all samples: (79 field samples and 8 negative controls) provided by ohmx.bio dry lab report. The median read lengths was 762 bp.

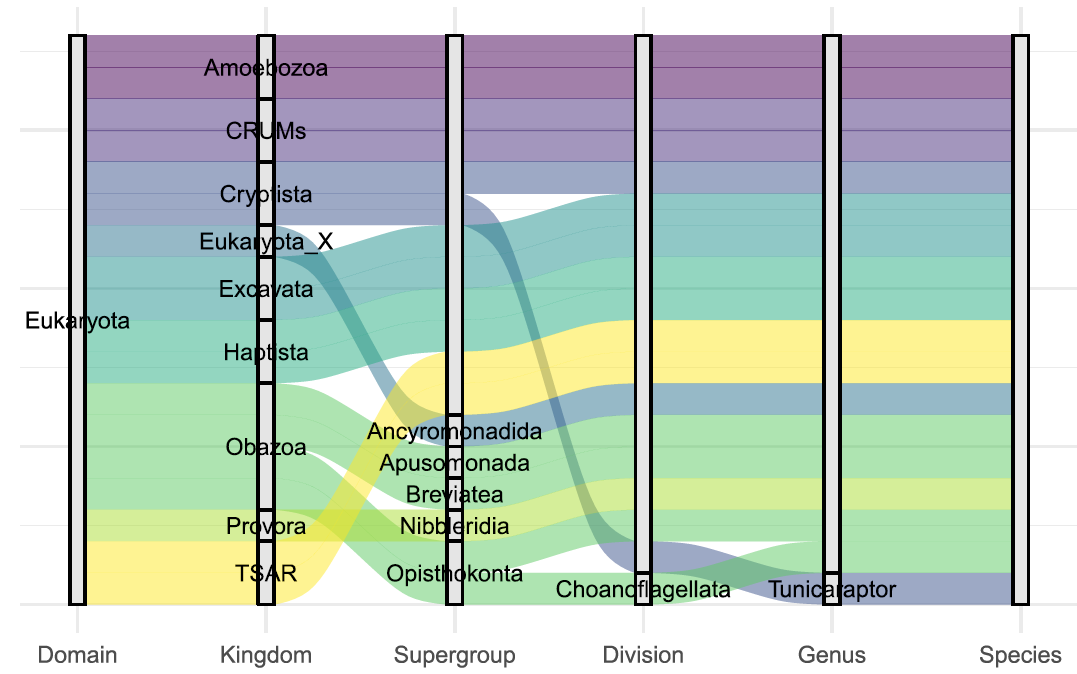

Figure S10. Alluvial plot illustrating the hierarchical taxonomic structure of eukaryotes retained for analysis focusing on protists and ranging from the domain to the species level.

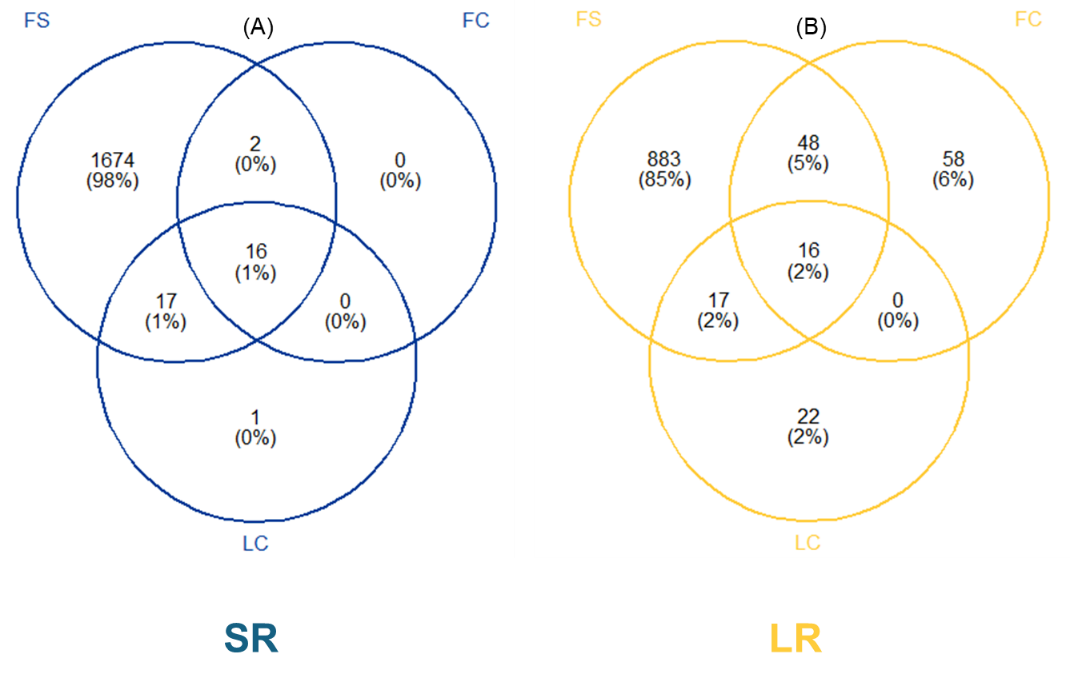

Figure S11. Venn diagrams of the number of identified taxa from negative field controls (FC) and lab controls (FC) and field samples (FS) with SR (in blue) and LR (in yellow) metabarcoding approaches to assess potential contamination.

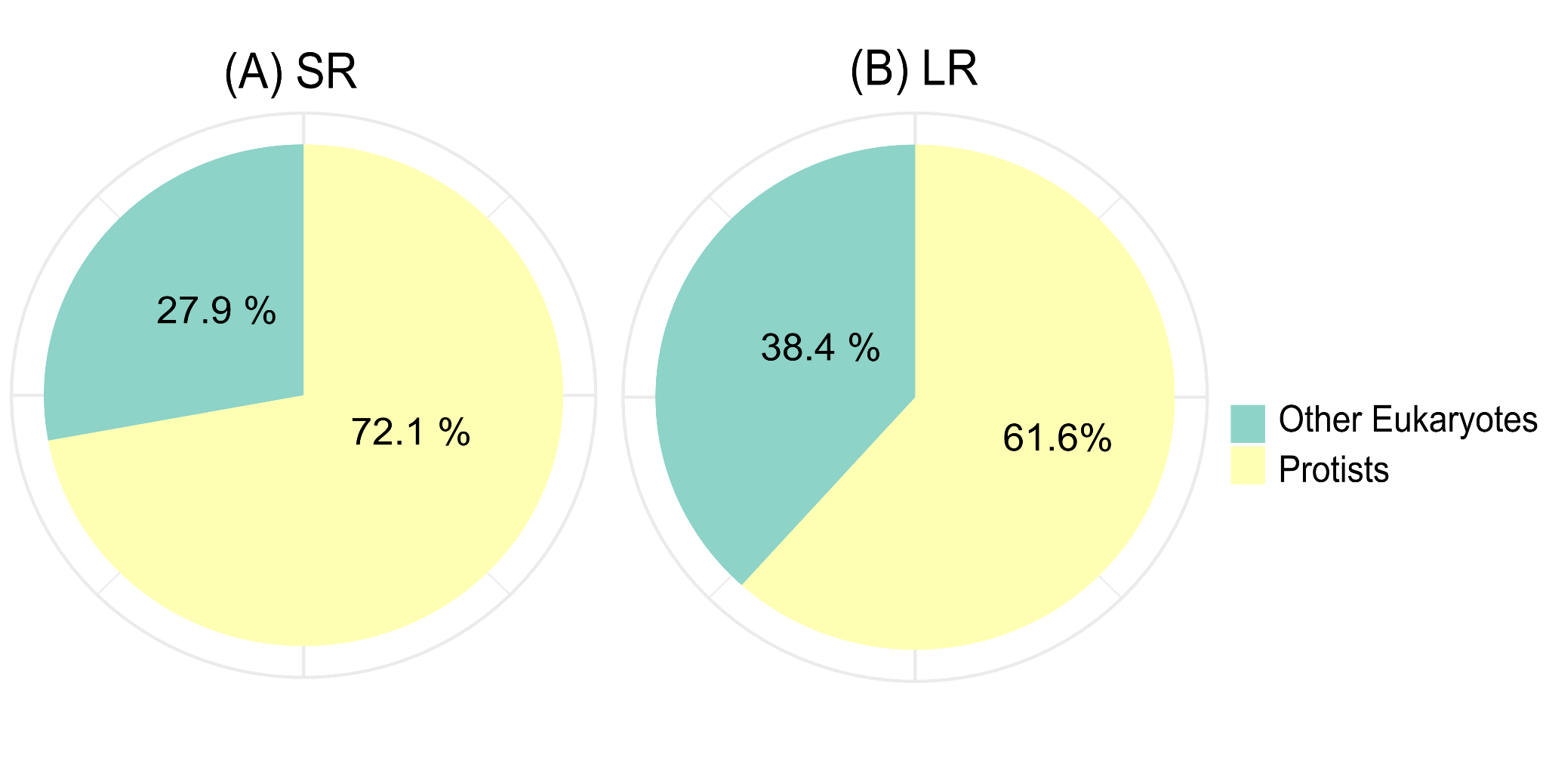

Figure S12. Pie charts representing the relative read abundance in percentages of protists as compared to other eukaryotes with (A) SR and (B) LR metabarcoding approaches for the entire dataset.

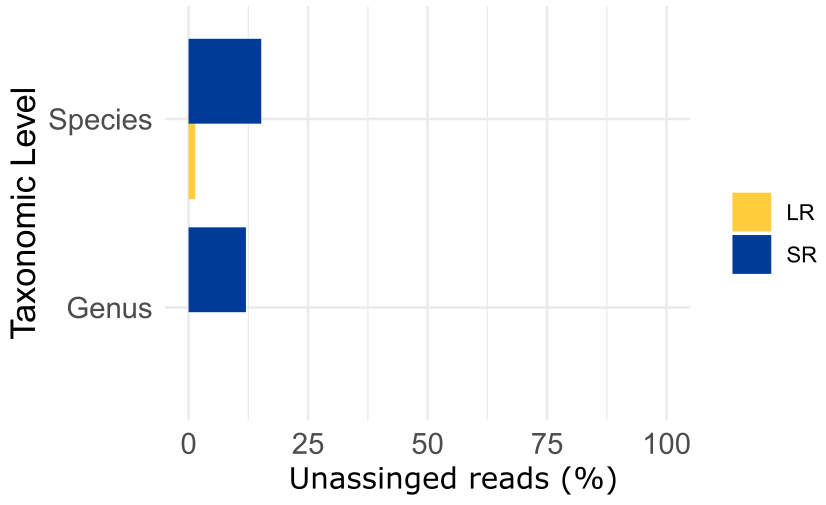

Figure S13. Horizontal histograms presenting the proportion of the relative abundance of unassigned reads in LR (yellow) and SR (blue) datasets at the genus and species level.

**Supplementary Table Legends**

Table S1. Station and sample description. Max. depth corresponds to the highest tide. SR and LR metabarcoding were conducted for a total of 79 field samples. Subsurface samples were collected at 1m depth, and epibenthic samples were collected 1m above the seabed. Note that MWO1 being closer to the coast was easier to sample under difficult weather conditions. See also Figure S2.

Table S2. Summary of protist taxa abundance according to three categories; *Abundant* (> 0.1%), *Intermediate Rare* (0.001 — 0.1 %), and *Rare* (< 0.001%); in the short-read (SR, yellow) and long-read (LR, blue) datasets.

Table S3. (A) List of Abundant Protist Genera (relative read abundance >0.1%), and unique detections showed in this study for both SR and LR metabarcoding approaches. Blue indicates the *Shared Abundant* genera (>0.1) detected in both SR and LR datasets (e.g., *Actinocyclus* detected in high abundance with both approaches). Pink refers to *Common* genera detected in both SR and LR but with contrasting abundance, for example the genus *Amphorellopsis* has a high abundance (>0.1%) in only one dataset (not specified for ease of reading), whilst the same genus is detected at lower abundance (<0.1%) in the other dataset. Grey corresponds to *Unique detection*, wherein an abundant genus is only detected by one approach (i.e., either SR or LR metabarcoding) whilst the genus is not detected by the other approach, which was the case for *Emiliana* (Haptophyta) and *Bellerochea* (Bacillariophyceae).

Table S4. Summary of the short-read metabarcoding dataset, including the number of total reads (non-rarefied samples) and alpha diversity estimates (rarefied samples). When applicable, maximum values are formatted in **bold green**, and minimum values in **bold red**.

Table S5. Summary of the short-read (**A**) and the long-read (**B**) metabarcoding datasets, including numbers of total reads (non-rarefied samples) and alpha diversity estimates (rarefied samples). When applicable, maximum values are formatted in **bold green**, and minimum values in **bold red**. For easy reading, samples are sorted according to locations (i.e., MOW1, WO5, WO8).

Table S6. Summary table of minimum, maximum and average values of alpha diversity estimates, for short-read (SR) metabarcoding (A) and long-read (LR) metabarcoding (B).

Table S7. Summary of the results of statistical tests for relevant factors using Kruskal-Wallis and Wilcoxon. Significant differences are highlighted in **bold** when applicable.
